# Unravelling the anti-cancer mechanisms elicited by non-covalent thioredoxin reductase inhibitors for triple negative breast cancer therapy

**DOI:** 10.1101/2025.09.04.674327

**Authors:** Abigail Rullo, Brenna Flowers, Keacha Chang, An Zhang, Valentina Z. Petukhova, Luke Harding, Sammy Y. Aboagye, Maurizio Bocchetta, Wei Qiu, David L. Williams, Francesco Angelucci, Pavel A. Petukhov, Irida Kastrati

## Abstract

Thioredoxin reductases (cytosolic TXNRD1 and mitochondrial TXNRD2) are antioxidant enzymes often overexpressed in tumors, including triple negative breast cancer (TNBC), making them promising targets for cancer therapy. Inhibiting these enzymes may worsen the already elevated oxidative stress in cancer cells, ultimately leading to cell death through a pro-oxidant mechanism. However, selectively targeting TXNRDs has been challenging due to the traditional reliance on covalent inhibition strategies. Recent studies have identified a druggable allosteric pocket in this enzyme family, paving the way for the development of novel non-covalent inhibitors, referred to as TXNRD(i)s. These inhibitors have been tested in TNBC models and have demonstrated a range of anti-cancer effects.

To understand the molecular and cellular consequences of TXNRD(i)s, we conducted unbiased transcriptomic analyses and found that the gene expression changes induced by TXNRD(i) treatment closely mirror those resulting from TXNRD1 silencing, reinforcing TXNRD1 as the primary therapeutic target. While TXNRD(i) treatment increases redox stress in TNBC cells, this is not the main driver of the anti-cancer effect. Instead, TXNRD(i)s potently inhibit cell proliferation and induce G1 phase cell cycle arrest. Notably, supplementing cells with exogenous deoxynucleotides restores cell viability, cell cycle progression and partially reverses cell death. These findings indicate that TXNRD(i)s deplete endogenous deoxynucleotide pools and impair ribonucleotide reductase activity as the main mechanism of anti-cancer effects. We further demonstrate that TXNRD(i)s inhibit both TXNRD1 and TXNRD2, and that dual inhibition is more effective in suppressing TNBC cell growth. *In vivo*, TXNRD(i) treatment significantly impairs TNBC xenograft tumor growth and reduces proliferation-related genes. Collectively, these findings challenge the prevailing paradigm that all TXNRD inhibitors function through a pro-oxidant mechanism, instead highlighting that non-covalent TXNRD(i)s exert their effects by blocking proliferation offering a compelling therapeutic strategy for TNBC and potentially other cancers with elevated TXNRD expression.

**Highlights:**

Non-covalent TXNRD inhibitors increase intracellular redox stress, but this is not the main driver of anti-cancer effects
TXNRD(i)s inhibit both cytosolic TXNRD1 and mitochondrial TXNRD2 enzymes
Both TXNRD1 and TXNRD2 are required for growth and proliferation of triple negative breast cancer cells
Halted proliferation through ribonucleotide reductase dysfunction emerges as the primary driver of anti-cancer effect

**Graphical Abstract:** 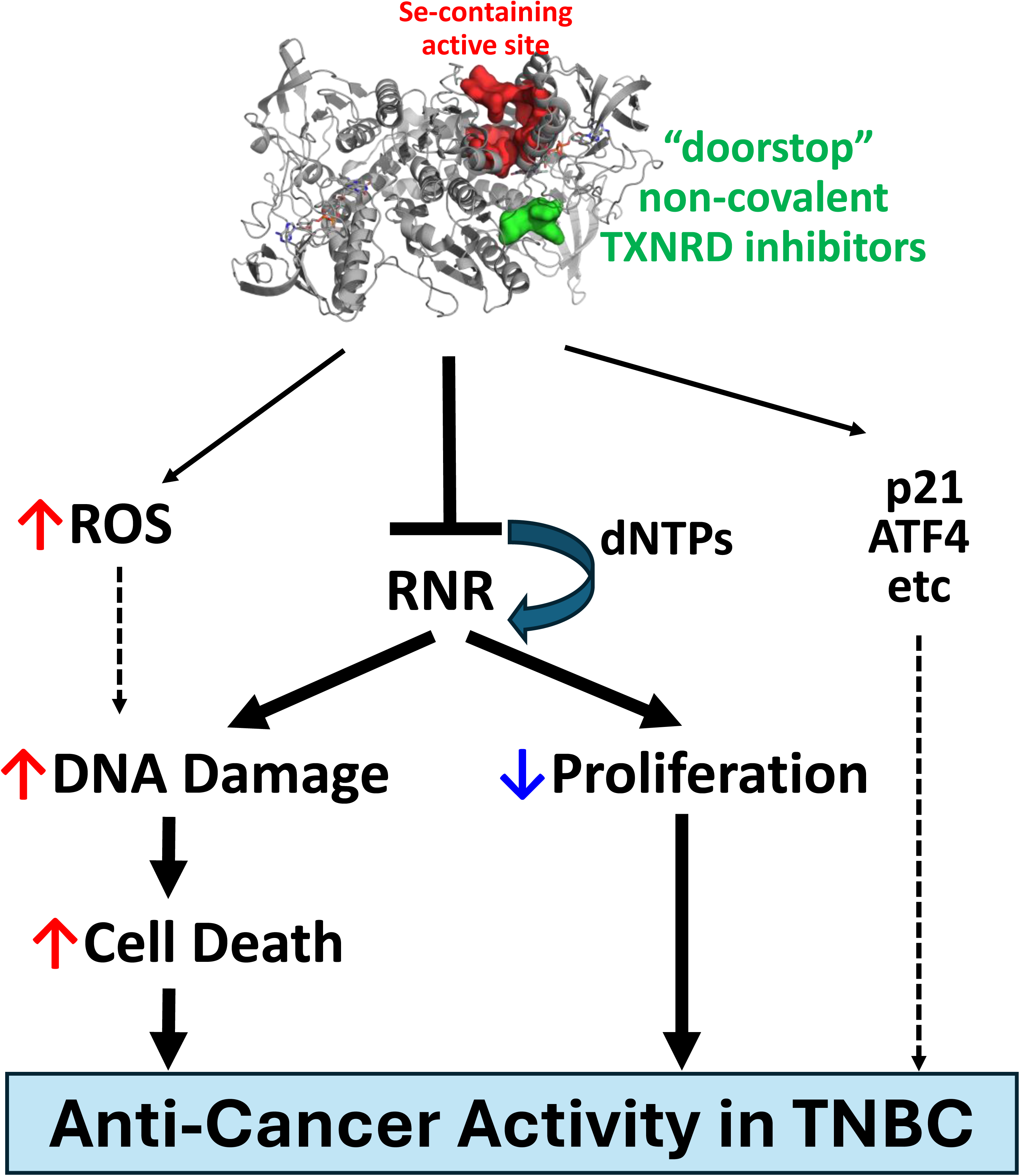

## 1. Introduction

The idea of targeting redox homeostasis as an anti-cancer strategy is not new. However, despite extensive research, translating this concept into effective treatments remains challenging. Redox homeostasis refers to the balance between reactive oxygen species (ROS) and the cell’s antioxidant defense mechanisms. ROS influence every cancer hallmark, including DNA damage and genetic alterations, cell death, metabolic rewiring, therapy resistance, escape from tumor immune microenvironment, and metastasis [1, 2]. In cancer cells low to moderate ROS levels induce genetic changes that are essential for cancer initiation, proliferation, and progression, as well as the development of therapeutic resistance. This positions the thioredoxin (Trx) and the glutathione (GSH) antioxidant systems as pivotal regulators of redox homeostasis and oncogenesis.

In breast cancer, studies using murine mammary gland tumor models have shown that GSH is required for tumor initiation [3]. However, once tumors are established, inhibiting GSH alone is ineffective due to compensatory upregulation of the Trx pathway [3]. In contrast, targeting the Trx pathway has demonstrated therapeutic benefits in limiting tumor progression [3]. The Trx system depends on the ubiquitously expressed selenoproteins TXNRD1 (cytosolic) and TXNRD2 (mitochondrial) isoenzymes along with NAPDH to maintain reduced Trx levels (Trx1 in cytosol; Trx2 in mitochondria) and support essential housekeeping and antioxidant defenses [4–6]. Our findings further underscore the importance of the Trx-TXNRD axis, showing that TXNRDs’ expression is elevated in triple-negative breast cancer (TNBC) and correlates with poor patient outcomes. TNBC is a heterogeneous, highly proliferative breast cancer subtype characterized by a high risk of recurrence and metastasis, the absence of targeted therapies, and the poorest prognosis among breast cancer types. Given the urgent need for new treatment strategies, targeting the Trx-TXNRD pathway offers a promising therapeutic approach for TNBC.

TXNRD enzymes reduce the oxidized disulfide-form to regenerate the catalytic dithiol of Trx, their main endogenous substrate. This reduced Trx, in turn, supports the activity of multiple downstream targets, including peroxiredoxins and other protein substrates. TXNRDs can also reduce a variety of other thioredoxin-fold proteins, such as protein disulfide-isomerase (PDI), as well as small-molecule substrates like menadione and lipoic acid (reviewed in [7]). Reduced Trx interacts with different downstream factors that play central roles in apoptosis, survival and gene transcription; all processes especially relevant to cancer. These include nuclear factor κB (NFκB), p53, apoptosis signal-regulating kinase (ASK1) and nuclear factor erythroid 2-related factor 2 (NRF2), among others [6]. Critically, Trx also supports the function of ribonucleotide reductase (RNR), the enzyme responsible for the rate-limiting step in the conversion of ribonucleotides (NTPs) into deoxyribonucleotides (dNTPs), which are required for DNA replication and repair [8]. Trx was originally discovered by Peter Reichard and colleagues in 1963 in the context of RNR activity in *E.coli*, where it serves as its reducing agent [9]. Most of our understanding of RNR regulation by the Trx system comes from studies in *E.coli* and yeast, with relatively limited insights from mammalian systems. RNR is a heterotetramer composed of two homodimeric subunits: RRM1, which contains the catalytic site, and RRM2, which contains a diferric iron center essential for generating the tyrosyl radical required for RRM1 activity [8]. Notably, a recent study found that depletion of Trx1 or TXNRD1 in lung cancer cells was synthetically lethal when combined with CHK1 inhibition, due to disrupted RNR function involving both RRM1 and RRM2 [10].

Despite its therapeutic potential, clinical progress in targeting TXNRDs has been hindered by the lack of selective and safe inhibitors. Traditional TXNRD inhibitors are covalent and/or irreversible, often interacting with off-target proteins and macromolecules, resulting in significant toxicity (reviewed in [11]). The use of these inhibitors also complicates the interpretation of biological outcomes. We recently discovered a new class of non-covalent TXNRD inhibitors (TXNRD(i)s) that bind to a newly identified allosteric regulatory site on the enzyme, termed the “doorstop pocket” [11, 12]. These novel inhibitors show potent anti-cancer activity in TNBC cells by reducing cell viability, increasing cell death, impairing invasion, and suppressing both clonogenic and mammosphere formation, while sparing normal breast epithelial cells [13]. Importantly, these compounds demonstrated specific inhibition of TXNRD activity both *in vitro* and *in vivo*, providing strong evidence for a defined on-target molecular mechanism of action. In preclinical TNBC models, two of these inhibitors, 8VP101 and 9VP19, significantly suppressed tumor growth, offering compelling proof-of-concept and validating the therapeutic promise of this pharmacologic strategy [13]. The pleiotropic anti-cancer effects observed with these non-covalent TXNRD(i)s underscore the critical role of TXNRD enzymes in driving oncogenic processes. These findings prompted us to further investigate the cellular mechanisms underlying their therapeutic activity.

## 2. Results

### 2.1. The transcriptome regulated by non-covalent TXNRD(i)s’ in TNBC largely overlaps with silencing of TXNRD1

The first-in-class non-covalent TXNRD(i)s, 8VP101 and 9VP19, were designed to bind to an allosteric regulatory site within the enzyme disrupting the electron flow from NADPH to reduce the disulfide of Trx and other substrates [11, 12]. As TXNRDs are selenoenzymes, the addition of exogenous selenium is expected to mitigate inhibition, potentially by increasing TXNRD abundance or activity [14]. Consistent with this notion, 8VP101, previously characterized in TNBC models, demonstrated diminished inhibitory effects in the presence of selenium (Supp. Fig. 1A). To investigate the global transcriptomic changes induced by TXNRD inhibition, RNA sequencing was performed on MDA-MB-231 TNBC cells treated with 10 µM of either 8VP101 or 9VP19 for 24 hours, with vehicle-treated cells as controls. TXNRD1 is regarded as the primary therapeutic target and non-covalent TXNRD(i)s effectively suppress TXNRD1 activity in TNBC cells [13]. Concurrently, RNA sequencing was conducted in cells transfected with either siTXNRD1 or non-silencing control siNeg for 48 hours (Supp. Fig. 1B). Each experimental condition was assessed in triplicate biological replicates (Supp. Fig. 1C-D), and hierarchical clustering of the resulting transcriptomes is presented in Supp. Fig. 1E. A Venn diagram (Fig. 1A) illustrates that the majority of transcripts altered by TXNRD1 silencing or pharmacologic inhibition via TXNRD(i)s are shared. However, the distinct pools of significantly altered transcripts (log2FC ≥ 0, *p*-value ≤ 0.05) depicted in a table shown in Fig. 1B reveal that the transcriptomic impact of TXNRD(i)s is approximately threefold greater than that of siTXNRD1. Comprehensive gene enrichment and pathway analyses were then applied to the significantly regulated transcriptomes.

**Figure 1.**
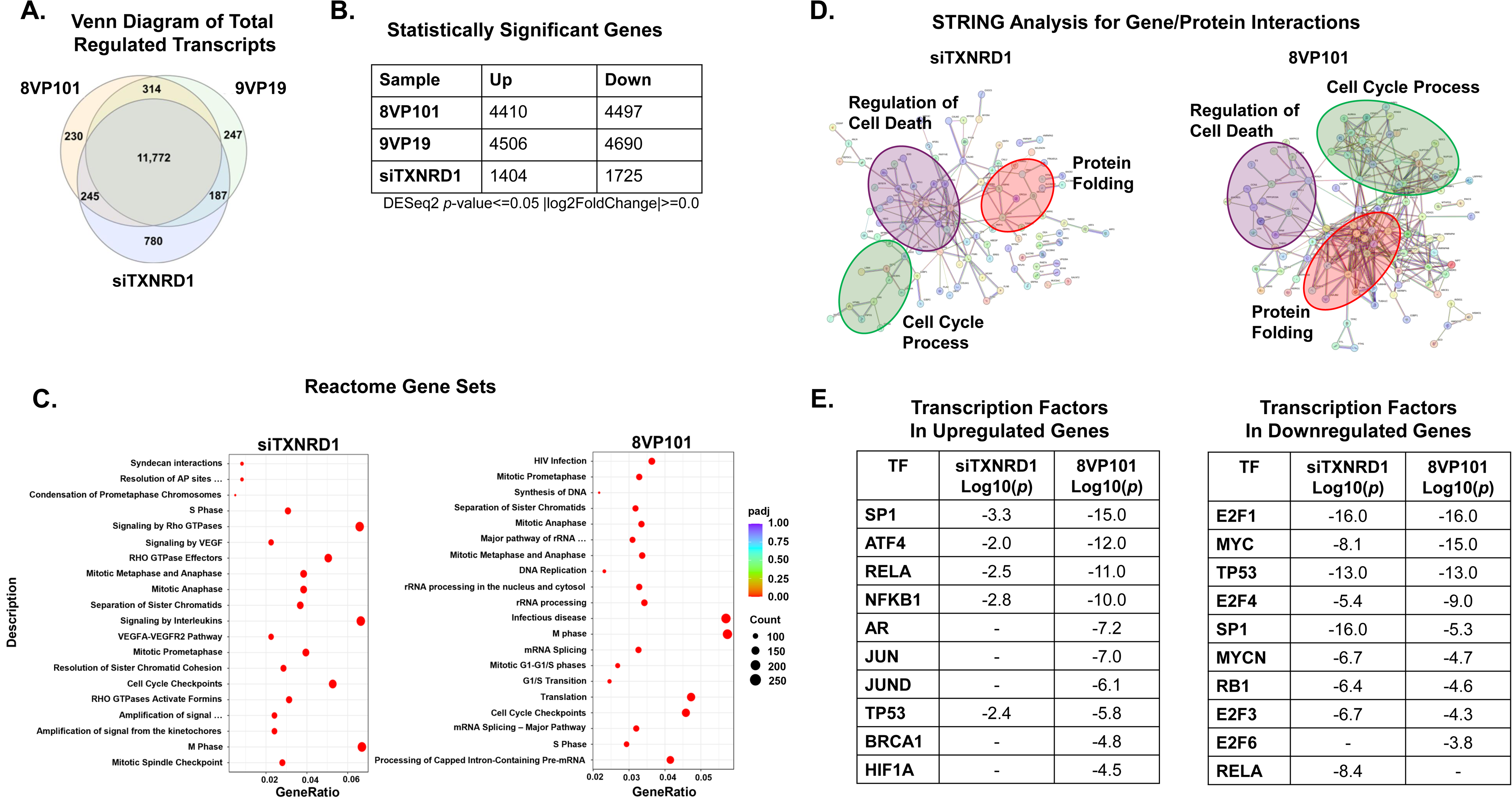
TXNRD(i) regulate unique gene signatures that largely overlap with silencing of TXNRD1. A. Venn diagram showing the overlap of the total differentially regulated transcripts between TXNRD(i)-treated samples (8VP101 and 9VP19, 10µM each) and siTXNRD1-transfected sample in MDA-MB-231 cells. 8VP101 and 9VP19 samples were compared to the vehicle-treated control, while siTXNRD1-transfected samples were compared to the siNEG-transfected control. B. Significantly regulated transcripts identified using the cut offs p-value<=0.05 and Log2FC>=0.0. 8VP101 and 9VP19 samples were compared to the vehicle-treated control, while siTXNRD1-transfected samples were compared to the siNEG-transfected control. C. Reactome enrichment analysis scatter plot. Gene ratio indicates the ratio of the number of differentially expressed genes within the reactome ID (identified by the pathway description) to the number of differentially expressed genes within the reactome database. Count indicates the number of genes within the indicated reactome ID. D. String analysis of the top 150 genes regulated by siTXNRD1 transfection or 8VP101 treatment compared to siNEG and vehicle, respectively. siTXNRD1: Regulation of Cell Death FDR: 0.0430, Protein Folding FDR: 0.0077, Cell Cycle Process FDR: 0.0279. 8VP101: Regulation of Cell Death 0.00031, Cell Cycle Process FDR: 8.56e-05, Protein Folding FDR: 7.01e-07. E. Metascape analysis identifying transcription factors regulating the top 2,500 upregulated genes (left) and top 2,500 downregulated genes (right), ranked by –log₁₀(p-value). (–) indicates not detected.

*Reactome* gene enrichment analysis reveals that cell cycle control and DNA replication processes are prominently represented in both siTXNRD1 and TXNRD(i)-treated cells. These include *M Phase*, *S Phase*, *G1/S Transition*, *Cell Cycle Checkpoints*, and *DNA Replication* signatures (Fig. 1C, Supp. Fig. 2A). STRING analysis further underscores the prominence of cell cycle processes, indicated by green bubbles in Fig. 1D and Supp 2B, which are regulated by siTXNRD1, 8VP101 and 9VP19. Additionally, *Cell Cycle* processes are linked to *Protein Folding* (red circles) and *Regulation of Cell Death* (purple circles), both similarly influenced by siTXNRD1, 8VP101 (Fig. 1D) and 9VP19 (Supp. Fig. 2B). The enrichment of *Cell Death* processes by 8VP101 and 9VP19 aligns with our previous findings [13]. Likewise, *Metascape* analysis highlights the *Mitotic Cell Cycle*, *DNA Metabolic Process* and *Regulation of Cell Cycle* as the top downregulated processes (Supp. Fig. 3A), whereas *Response to Oxidative Stress* and *Cellular Catabolic Process* as the top upregulated processes (Supp. Fig. 3B). No distinct pathway is identified within the smaller, unique sets of transcripts regulated by either TXNRD(i)s or siTXNRD1 alone (Supp. Fig. 4). The top transcription factors implicated in mediating the transcriptional changes with siTXNRD1 or TXNRD(i)s include SP1, E2F1, ATF4, MYC, RELA and TP53 (Fig. 1E and Supp. Fig. 2C), aligning with the fact that the Trx-TXNRD axis supports transcription factor such as NFκB and p53. Of note, ATF4, a transcription factor of the Unfolded Protein Response (UPR), is consistent with *Protein Folding* process noted earlier and further substantiated by significant upregulation of UPR genes such as, PERK, CHOP and BIP (Supp. Fig. 5B). E2F1, specifically linked to cell cycle regulation, is investigated again in the subsequent sections. Given the similarities in the transcriptome regulated by either of TXNRD(i)s, 8VP101 or 9VP19, an overlapping signature is used and referred to as *TXNRD(i) signature* in the rest of the text.

### 2.2. TXNRD(i) treatment increases redox stress, but it is not critical to cytotoxicity

*Oxidative Stress Response* emerged as a prominent pathway in siTXNRD1-transfected cells and was similarly enriched in TNBC cells treated with TXNRD(i)s (Fig. 2A) consistent with inhibition of the antioxidant thioredoxin system, which mitigates ROS and peroxides via electron donation to peroxiredoxins. To assess intracellular ROS levels, we utilized the fluorescent, cell-permeable probe H2-DCFDA and analyzed cells via flow cytometry after treatment with 10 µM 8VP101 for 2, 6, 18, and 24 hours. Following treatment, cells were incubated with 1 µM H2-DCFDA for 15 minutes before FACS analysis, with the gating strategy outlined in Supp. Fig. 6A. This probe detects various ROS, including hydrogen peroxide, hydroxyl radicals, and peroxyl radicals [15]. We observed a time-dependent increase in intracellular ROS in TNBC cell lines MDA-MB-231 and HCC1806, as well as the non-tumorigenic breast epithelial line MCF-10A upon 8VP101 treatment, with ROS levels plateauing at approximately a two-fold increase across all lines (Fig. 2B). However, despite similar ROS induction, cell viability varied significantly, with TNBC cells displaying greater sensitivity to inhibition by 8VP101 compared to MCF-10A (Fig. 2C).

**Figure 2.**
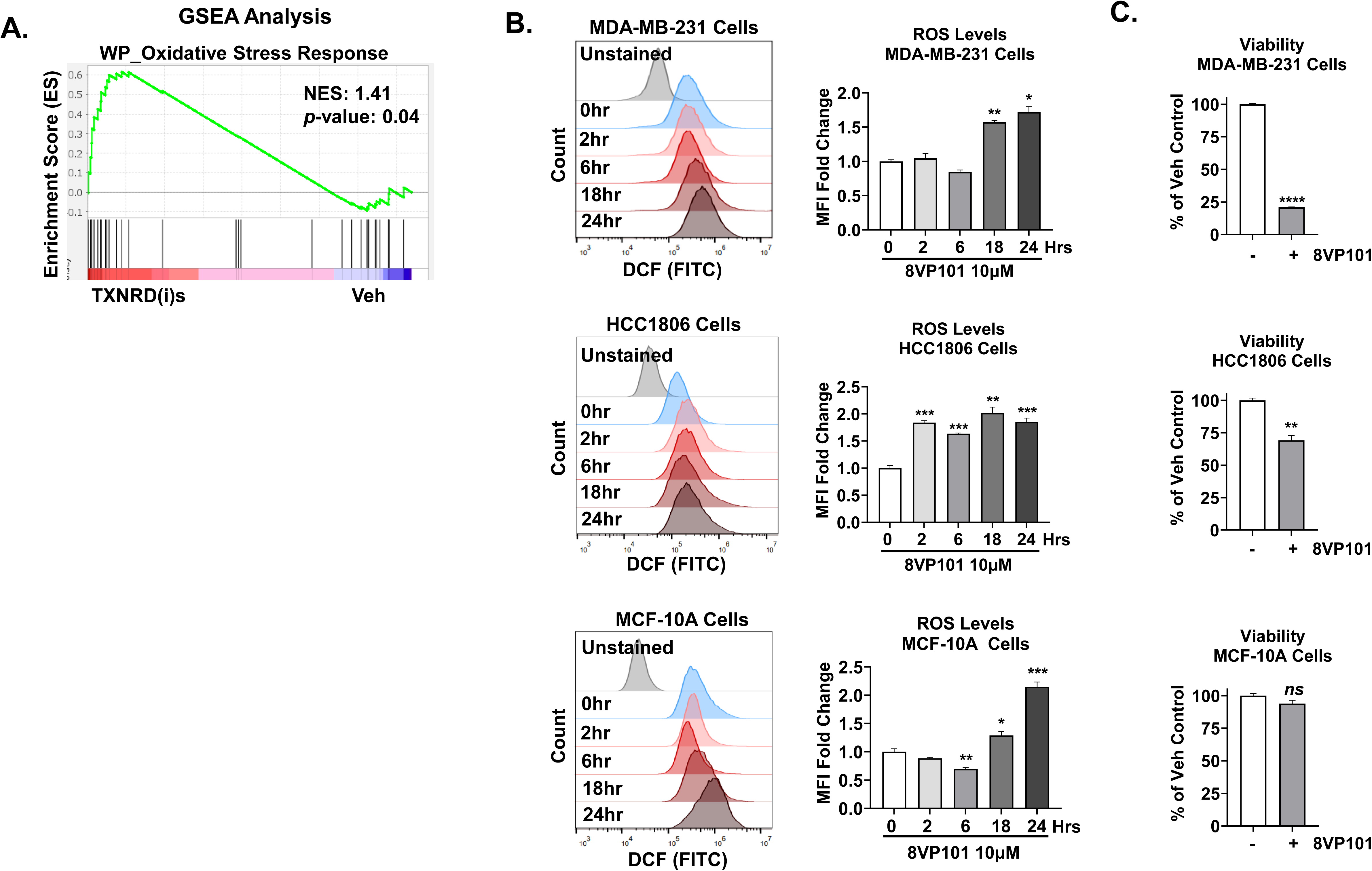
TXNRD(i)s increase ROS levels which do not correlate with cell viability. A. Gene set enrichment analysis of *Oxidative Stress Response* gene set by WikiPathways. NES and nominal p-value were determined by the GSEA software. B. ROS quantification using H_2_DCFDA by flow cytometry of MDA-MB-231 (top) (n=2), HCC1806 (middle) (n=3), and MCF-10A (bottom) (n=3) cells. Cells were treated with 10µM 8VP101 for 0, 2, 6,18, and 24 hours, harvested and incubated with 1µM H_2_DCFDA for 15 minutes. Representative histograms (left) of measured DCF mean fluorescence intensity (MFI). Statistical significance was determined by unpaired t-test relative to 0-hour time point. C. Cell Viability determined by crystal violet. MDA-MB-231 (top), HCC1806 (middle), and MCF-10A (bottom) cells treated with 10µM 8VP101 for 24 hours. Cells were fixed and stained with crystal violet, then solubilized and absorption values are normalized to vehicle control set as 100%. Statistical significance was determined by unpaired t-test. Data is presented as the mean +/-sem. *p≤0.05; **p≤0.005; ***p≤0.001; ****p≤0.0001; ns: not significant.

To assess the role of ROS induction in the anti-cancer effects of TXNRD(i)s, we employed two ROS scavengers: N-acetyl cysteine (NAC) and α-tocopherol (α-T, vitamin E). NAC replenishes glutathione (GSH), the most abundant cellular ROS detoxifier [16], and effectively scavenges hydrogen peroxide-induced redox stress in TNBC cells (Supp. Fig. 6B). α-T, a naturally occurring lipophilic antioxidant, quenches ROS and interrupts peroxidation chain reactions by neutralizing peroxyl radicals [17]. Treatment with either NAC or α-T effectively restored intracellular redox homeostasis to baseline levels in TXNRD(i)-treated cells, with α-T showing greater efficacy (Fig. 3A, 3C). However, despite mitigating ROS, neither NAC nor α-T rescued cell viability in MDA-MB-231 or HCC1806 cells treated with low concentrations of 8VP101 (Fig. 3B, 3D) or a high concentration of 8VP101 (Supp. Fig. 6C). These findings indicate that while TXNRD(i) treatment elevates redox stress, ROS induction alone is not the primary driver of the anti-cancer effects, as ROS scavenging does not reverse the cytotoxicity.

**Figure 3.**
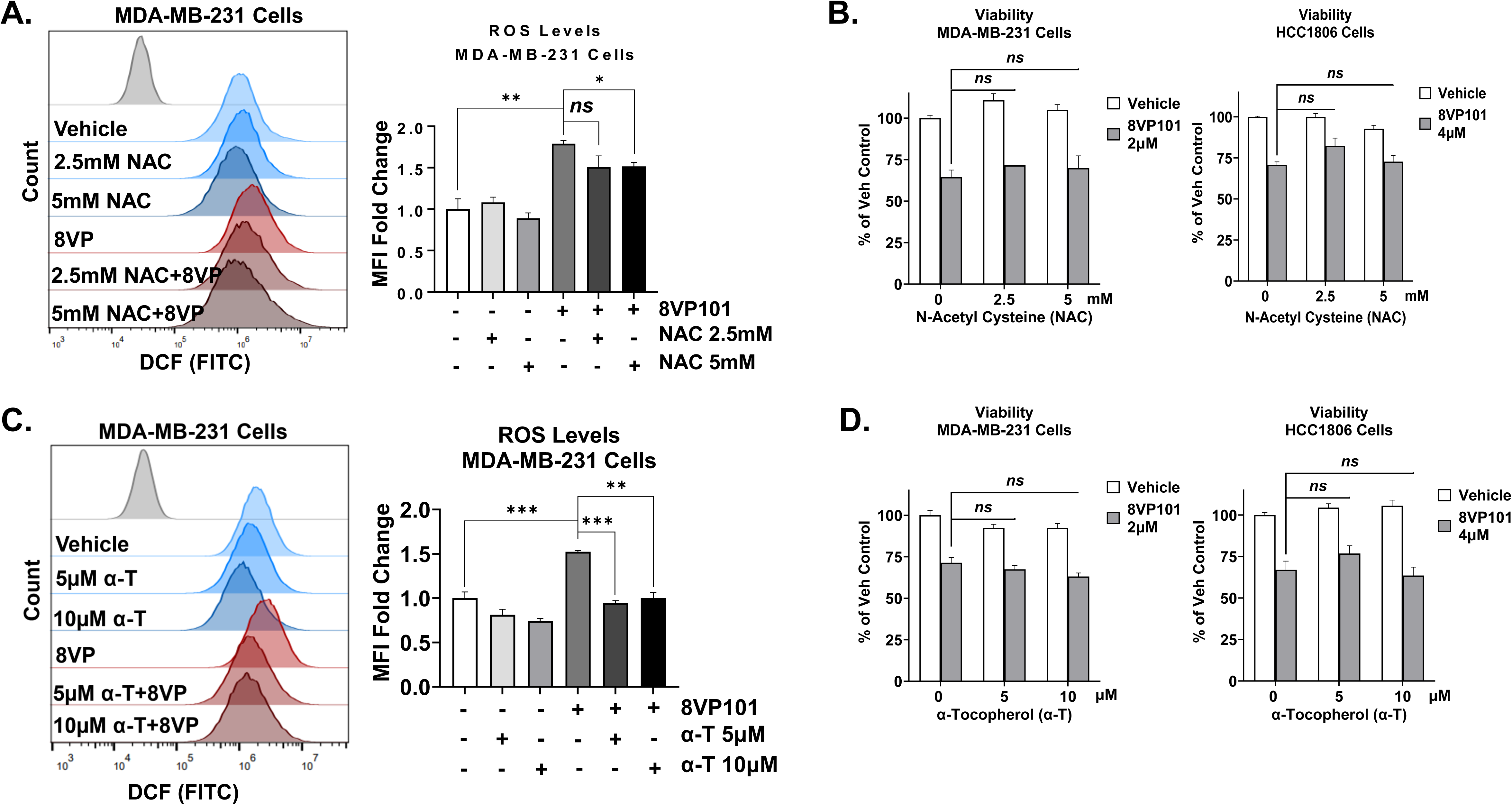
Antioxidants cannot rescue the TXNRD(i)s’ effect on cell viability. A. ROS quantification using H_2_DCFDA by flow cytometry of MDA-MB-231 cells. Cells were treated with 10µM 8VP101 and 2.5mM or 5mM NAC for 24 hours, harvested and incubated with 1µM H_2_DCFDA for 15 minutes. Representative histogram (left) of measured DCF MFI. Statistical significance was determined by unpaired t-test. B. Cell Viability determined by crystal violet. MDA-MB-231 (left) and HCC1806 (right) cells were treated with 2µM and 4µM 8VP101, respectively, and 2.5mM or 5mM NAC for 72 hours. Cells were fixed and stained with crystal violet. Statistical significance was determined by 1-way ANOVA followed by Tukey’s test. C. ROS quantification using H_2_DCFDA by flow cytometry of MDA-MB-231 cells. Cells were treated with 10µM 8VP101 and 5µM or 10µM of ⍺-tocopherol (⍺-T) for 24 hours, harvested and incubated with 1µM H_2_DCFDA for 15 minutes. Representative histogram (left) of measured DCF MFI. Statistical significance was determined by unpaired t-test. D. Cell Viability determined by crystal violet. MDA-MB-231 (left) and HCC1806 (right) cells were treated with 2µM and 4µM 8VP101, respectively, and 5µM or 10µM ⍺-Tocopherol (⍺-T) for 72 hours. Cells were fixed and stained with crystal violet. Statistical significance was determined by 1-way ANOVA followed by Tukey’s test. Data is presented as the mean +/-sem. *p≤0.05; **p≤0.005; ***p≤0.001; ns: not significant.

### 2.3. TXNRD(i)s block proliferation, induce cell cycle arrest and trigger cell death by depleting dNTP pools and impairing RNR function

Given the prominent *DNA Replication* (Fig. 4A) and *Cell Cycle* signatures enriched in the transcriptomics data following TXNRD(i) treatments, we further investigated these pathways. We observe a marked reduction in DNA synthesis as evidenced by decreased incorporation of 5-ethynyl-2′-deoxyuridine (EdU), a thymidine nucleoside analog, into newly synthesized DNA in cells treated with increasing concentrations of 8VP101 (Fig. 4B) with incorporation levels dropping from 57% to 24%. A broad reduction in TNBC cell proliferation following 8VP101 treatment was further supported by gene expression data (Fig. 4C); both *Ki67*, a marker of proliferation, and *E2F1*, a transcription factor promoting cell cycle progression, were downregulated, while p21, a cell cycle inhibitor, was significantly upregulated consistent with changes observed at the protein level (Fig. 4E). Similarly, cell cycle analyses in MDA-MB-231 and HCC1806 cells (gating strategy in Supp. Fig. 6D), reveal a significant reduction in S phase and an accumulation in G1 phase (Fig. 4D). This profile contrasts with the cell cycle effects induced by AF (Supp. Fig. 6E). While the immortalized MCF-10A cells exhibited a similar trend, the magnitude of cell cycle disruption was notably less pronounced (Fig. 4D, bottom panels).

**Figure 4.**
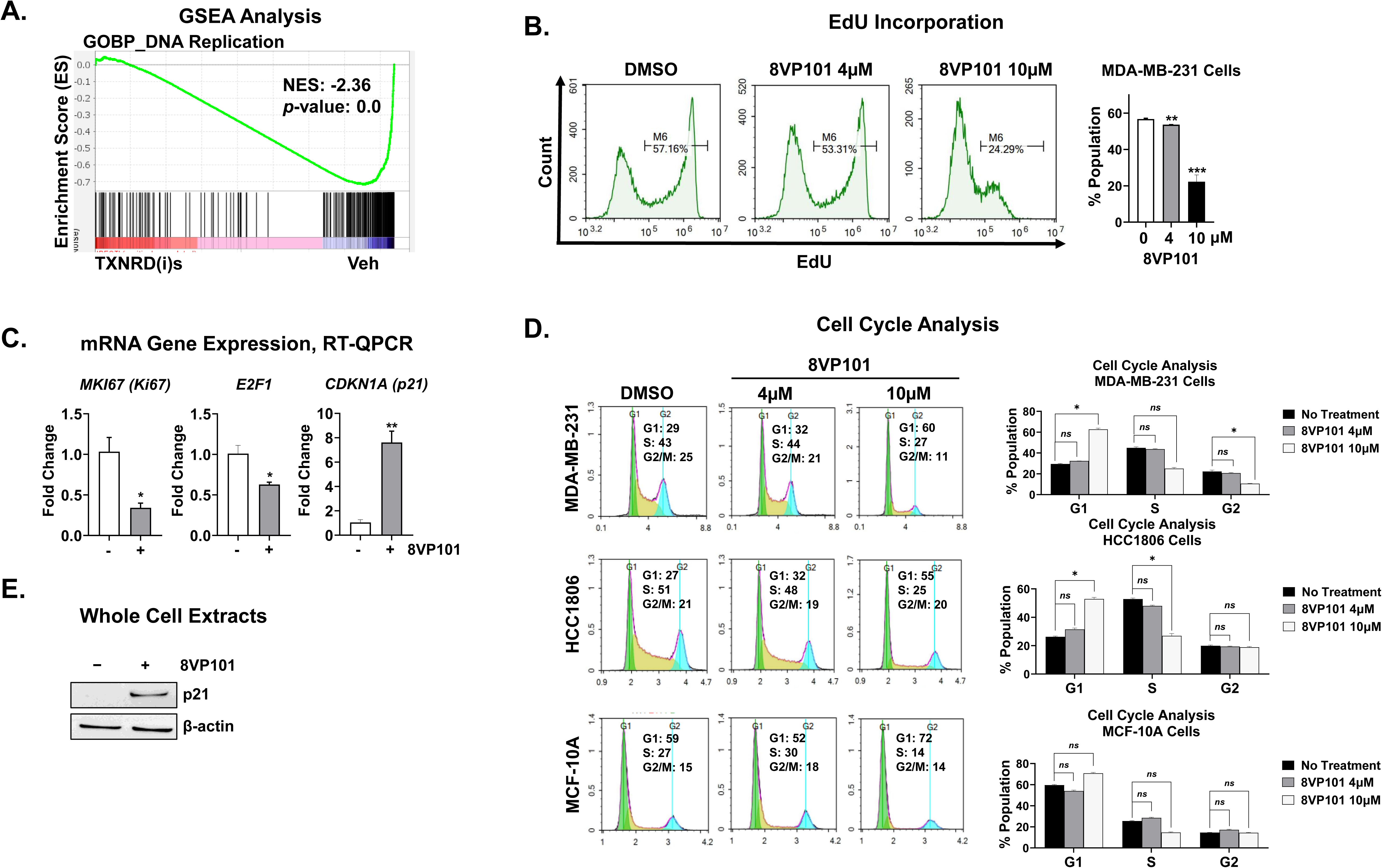
Treatment with TXNRD(i)s inhibits cells proliferation and disrupts cells cycle by inducing G1 arrest and reducing S phase. A. Gene set enrichment analysis of *DNA Replication* by Gene Ontology Biological Processes. NES and nominal p-value were determined by GSEA software. B. EdU incorporation assay by flow cytometry of MDA-MB-231 cells. Cells were treated with 4μM or 10μM 8VP101 for 24 hours and loaded with 1μM EdU for 3 hours before harvest and analysis. Representative histograms of EdU incorporation are shown on the left. Statistical significance was determined by unpaired t-test. C. Gene expression by RT-qPCR in MDA-MB-231 cells treated with 10µM 8VP101 for 24 hours. Relative gene expression is calculated using the ΔΔCt method normalized to β-actin. Statistical significance was determined by unpaired t-test. D. Cell Cycle Analysis using PI by flow cytometry of MDA-MB-231 (top), HCC1806 (middle), and MCF-10A (bottom) cells. Cells were treated with 4μM or 10μM 8VP101 for 24 hours. Statistical significance was determined by Kruskal Wallis test followed by Dunns test. E. Protein levels of p21 assessed by western blot. MDA-MB-231 cells were treated with 10μM 8VP101 for 6 hours. β-actin was used as the loading control. Data is presented as the mean +/-sem. *p≤0.05; **p≤0.005; ***p≤0.001; ns: not significant.

The Trx/TXNRD system is known to regulate DNA synthesis through redox control of ribonucleotide reductase (RNR), as demonstrated in studies using *E. coli* and yeast; however, evidence in mammalian systems remains limited. RNR catalyzes the reduction of ribonucleotides to deoxyribonucleotides (dNTPs), which are essential precursors for DNA synthesis and repair [8]. We observed a significant downregulation of *Purine and Pyrimidine Metabolism* signatures following TXNRD(i) treatment (Fig. 5A). Notably, the expression of RNR subunits RRM1 and RRM2 is significantly elevated in breast tumors compared to normal tissue, with a pronounced increase observed in TNBC tumors (Supp. Fig. 7A-B). Supplementation with exogenous dNTPs, the products of RNR activity, rescued both cell viability (Fig. 5B) and the proliferation arrest induced by TXNRD(i)s in the two TNBC cell lines by restoring progression through the G1 and S phases of the cell cycle (Fig. 5C, third and fourth bars). Given the role of dNTPs in DNA repair, we assessed DNA integrity using γH2AX as a marker of DNA damage [18]. TXNRD(i) treatment significantly increased γH2AX levels, which were partially reduced by dNTP supplementation (Fig. 5D). As DNA damage can lead to cell death, we examined this outcome and found that TXNRD(i)s increased both apoptosis and overall cell death (Fig. 5E, first and third flow panel), consistent with our prior findings [13]. Importantly, exogenous dNTPs significantly mitigated cell death (Fig 5E), indicating that impaired RNR function and subsequent dNTP depletion contributes to the cytotoxic effects of TXNRD(i)s.

**Figure 5.**
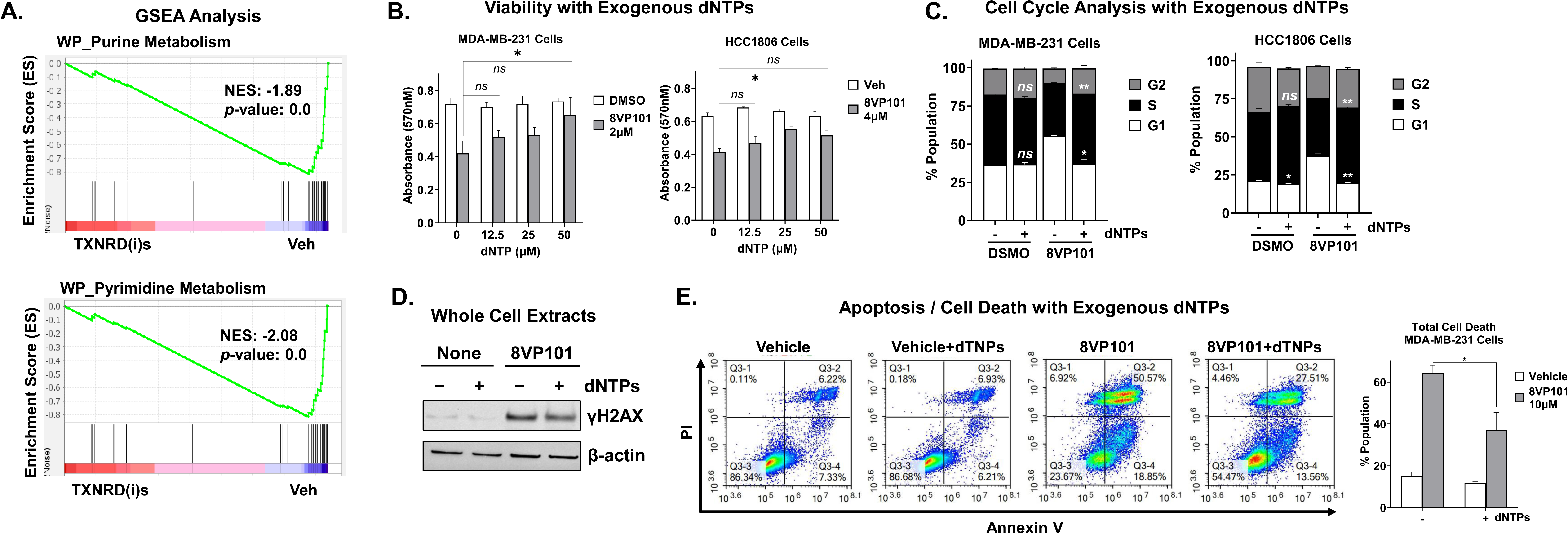
RNR dysfunction underlies the reduced cell viability, cell cycle arrest, and apoptosis induced by TXNRD(i)s. A. Gene set enrichment analysis of Purine Metabolism and Related Disorders and *Pyrimidine Metabolism* by WikiPathways. NES and nominal p-value were determined by GSEA software. B. Cell viability was determined by crystal violet. MDA-MB-231 (left) and HCC1806 (right) cells were treated with 2μM or 4μM 8VP101, respectively, and supplemented with 12.5μM, 25μM or 50μM exogenous dNTPs for 24 hours. Statistical significance was determined by 1-way ANOVA followed by Tukey’s test. C. Cell cycle analysis using PI by flow cytometry of MDA-MB-231 (left) and HCC1806 (right) cells. Cells were treated with 10μM 8VP101 and supplemented with 50μM exogenous dNTPs for 24 hours. Statistical significance was determined by Kruskal Wallis test followed by Dunns test between Veh and 8VP101 treated samples with or without exogenous dNTP supplementation. D. Protein levels of γH2AX assessed by western blot. MDA-MB-231 cells were treated with 10μM 8VP101 and supplemented with 50μM exogenous dNTPs for 6 hours. β-actin was used as the loading control. E. Cell death assessed by PI and annexin V staining of MDA-MB-231 cells measured by flow cytometry. Cells were treated with 10μM 8VP101 and supplemented with 50μM exogenous dNTPs for 24 hours. Statistical significance was determined by unpaired t-test. Data is presented as the mean +/-sem. *p≤0.05; **p≤0.005; ns: not significant.

Next, we evaluated the effect of 8VP101 in combination with standard-of-care therapies commonly used to treat TNBC, including the cytotoxic chemotherapies paclitaxel, doxorubicin, and cisplatin, as well as the poly ADP-ribose polymerase (PARP) inhibitor olaparib. Synergy scores were calculated using SynergyFinder, with scores ≥10 indicating synergy and scores between 1–10 considered additive [19]. While the three cytotoxic agents showed additive effects with synergy scores just below 10, olaparib demonstrated a synergy score >10, indicating a synergistic interaction with TXNRD(i) in suppressing TNBC cell growth with (Supp. Fig. 7C).

To assess whether TXNRD(i)s inhibit tumor growth *in vivo* by blocking proliferation, xenograft tumors were established in immunocompromised mice through the injection of human TNBC cells, MDA-MB-231 and HCC1806. Once tumors were established, mice were randomized to receive either 8VP101, 50 mg/kg, daily IP injection, or vehicle control, following the treatment scheme outlined in Fig. 6A. After three consecutive days of treatment, tumors were excised and analyzed for proliferation markers. We previously reported that 8VP101 significantly suppressed MDA-MB-231 tumor growth [13]. Similarly, we observed comparable suppression and even regression of HCC1806 xenograft tumor growth (Fig. 6B). Additionally, despite the brief treatment duration, proliferation markers were reduced (Fig. 6C). Collectively, these findings support halted proliferation via RNR dysfunction as the primary mechanism of action for non-covalent TXNRD(i)s in TNBC.

**Figure 6.**
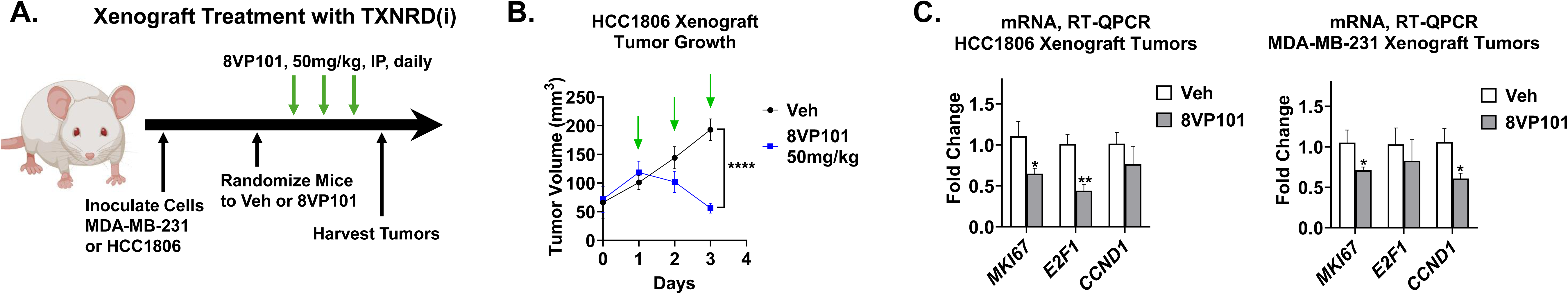
TXNRD(i) inhibits proliferation and growth of xenograft tumors. (A) HCC1806 cells were injected into bilateral mammary fat pads of 5-week old female athymic nude mice (n = 2-3 mice; n =4-6 tumors per group). Once tumors were consistently palpable, mice were randomized into vehicle or 50mg/kg 8VP101 groups and received daily intraperitoneal (IP) injections of the designated treatment daily. (B) HCC1806 xenograft tumors from schematic (A) were measured with digital calipers daily. Green arrows indicate treatment with TXNRD(i). (C) RNA was isolated from HCC1806 (n = 2 for vehicle group and 3 for 8VP101 group) and MDA-MB-231 (n = 2 for vehicle group and 3 for 8VP101 group) xenograft tumors from 6-week-old athymic nude female mice treated with vehicle or TXNRD(i)s for 3 consecutive days, then used for RT-qPCR for MKi67, E2F1, and CCND1. P-values show significance compared to untreated mice. Data is presented as the mean +/-sem. *p≤0.05; **p≤0.005; ****p≤0.0001.

### 2.4. TXNRD(i)s engage and inhibit both cytosolic and mitochondrial TXNRD enzymes for effective and broad anti-cancer effects

Given the pronounced anti-cancer effects of TXNRD(i)s and the three-fold larger transcriptomic response regulated by TXNRD(i)s compared to siTXNRD1 (Fig. 1B), we investigated whether TXNRD(i)s may have additional targets. The *doorstop pocket* of TXNRD1 is predicted to be conserved in other members of the thioredoxin reductase family, including the ubiquitously expressed mitochondrial TXNRD2 [11]. We employed the highly selective off–on fluorescent probe TRFS-green to measure TXNRD activity in live TNBC cells, as previously described [13, 20]. To distinguish between cytosolic and mitochondrial TXNRD pools, we performed simultaneous labeling with MitoTracker-red, which marks active mitochondria [21]. Co-localization of TRFS-green and MitoTracker-red signals appearing as orange enables quantification of mitochondrial TXNRD activity attributed to TXNRD2, whereas TRFS-green signal outside mitochondria reflects cytosolic TXNRD1 activity (representative images shown in Fig. 7A and Supp. Fig. 8A). Using this dual-labeling method, we confirmed that AF inhibits both TXNRD1 and TXNRD2 (Supp. Fig. 8B), consistent with its classification as a pan-TXNRD inhibitor [22]. Applying the same approach to 8VP101 and 9VP19, we found that these TXNRD(i)s also effectively inhibit both cytosolic and mitochondrial pools of TXNRDs (Fig. 7A and Supp. Fig. 8C), confirming that they function as pan-TXNRD inhibitors.

**Figure 7.**
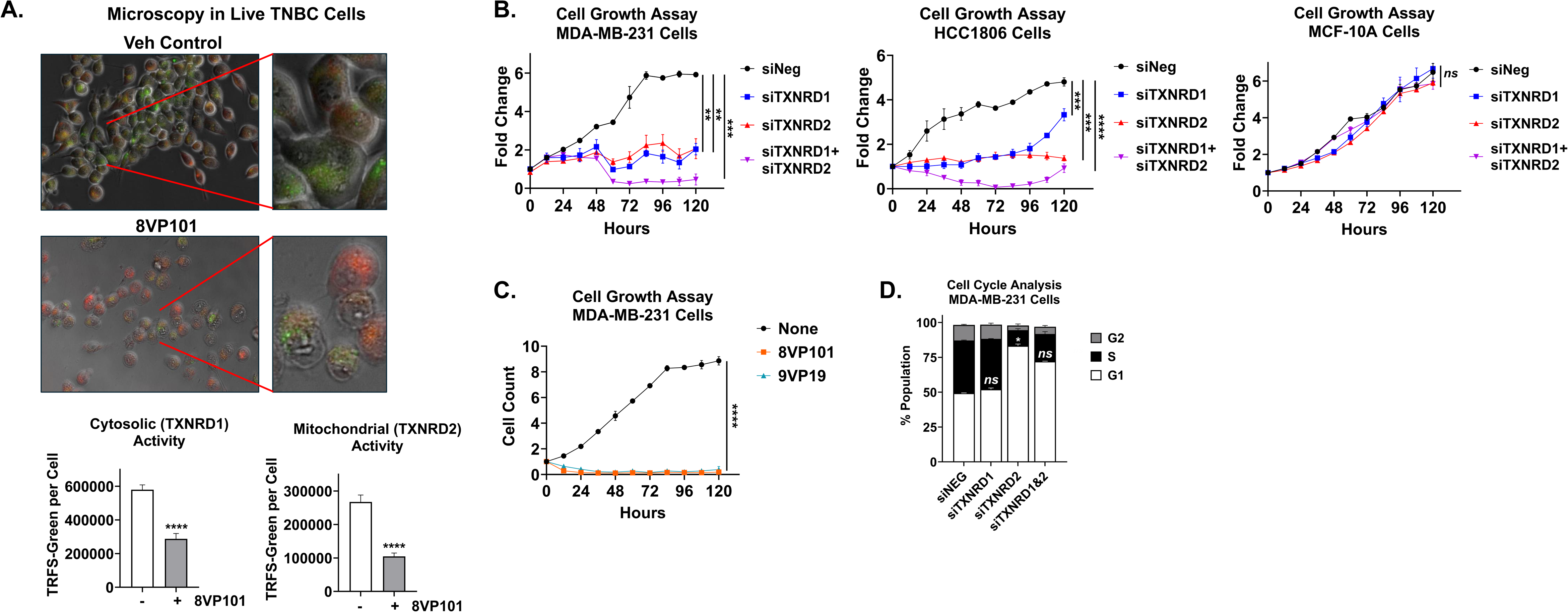
TXNRD(i)s inhibit TXNRD1 and TXNRD2 enzymes and both enzymes are critical for growth of TNBC cells. A. HCC1806 cells were pretreated with 8VP101 or vehicle control for 1 hour, then treated with 10µM of TRFS-green for 4 hours, followed by incubation with MitoTracker-red. Live cells were imaged on a Nikon Ti2E inverted microscope at 20x. Representative images shown. Cytosolic and mitochondrial TXNRD activity on average intensity per cell per field basis was quantified for 4µM 8VP101 treatment. Average intensity was calculated from n=3 biological replicates and n=9 technical replicates. ****p≤0.0001. B. MDA-MB-231, HCC1806 and MCF-10A cells were transfected with siNEG, siTXNRD1, siTXNRD2, or both siTXNRD1 and siTXNRD2, 10nM each, and monitored for cell growth every 12 hours over 120 hours using BioTek BioSpa. Data is shown as fold change normalized to initial timepoint. Growth curves are averages -/+ sem from n=3 biological replicates. End-point statistical significance was determined. **p≤0.005; ***p≤0.001; ****p≤0.0001. C. MDA-MB-231 cells were treated with DMSO, 8VP101 or 9VP19 at 10µM each and monitored for cells growth as in (B). End-point statistical significance was determined. ****p≤0.0001. D. Cell cycle analysis using PI by flow cytometry of MDA-MB-231 cells. Cells were transfected with 10nM siTXNRD1, 10nM siTXNRD2, or a combination of 10nM siTXNRD1 and 10nM siTXNRD2 for 24 hours. Statistical significance was determined by Kruskal Wallis test followed by Dunns test comparing siTXNRD1, siTXNRD2, and both siTXNRD1 and siTXNRD2 to siNEG control. Data is presented as the mean +/-sem. *p≤0.05; ns: not significant.

This finding prompted us to investigate the individual and combined roles of TXNRD1 and TXNRD2 in TNBC cells. We used small interfering RNA (siRNA)-mediated silencing and direct cell counting with the BioTek BioSpa system to assess cell growth in two TNBC cell lines, MDA-MB-231 and HCC1806. Silencing TXNRD1 alone (Fig. 7B, blue lines) resulted in significant growth suppression, although the effect diminished at later time points in HCC1806 cells, consistent with previous reports that identify TXNRD1 as the main cancer-relevant target [6], particularly in TNBC [13]. In contrast, silencing TXNRD2 (Fig. 7B, red lines) led to a significant and sustained reduction in cell growth over time in both cell lines, indicating that TXNRD2 is also required for TNBC proliferation and represents a previously underappreciated therapeutic target. Additionally, combined silencing of TXNRD1 and TXNRD2 (Fig. 7B, purple lines) produced a more pronounced effect, resulting in regression of the growth curves suggestive of cytotoxic, rather than merely cytostatic, effects. Notably, MCF-10A non-tumorigenic breast epithelial cells were unaffected by either individual or combined silencing (Fig. 7B, last panel), suggesting a potential therapeutic window. The cell cycle profiles following individual or dual TXNRD silencing closely mirrored those observed with TXNRD(i) treatment (Fig. 7). Similarly, the anti-proliferative effects of TXNRD(i)s in TNBC cells closely mimicked the effects of dual TXNRD1/2 silencing (Fig. 7C).

Together, these results support the conclusion that the enhanced anti-cancer activity of TXNRD(i)s arises from pan-TXNRD inhibition, which emerges as a more effective therapeutic strategy for TNBC. Other solid tumors show elevated expression of TXNRD1, TXNRD2 or both, including lung cancer (Supp. Fig. 9A). We evaluated and found that several lung cancer cell models express both TXNRD1 and TXNRD2. Furthermore, using two TXNRD(i)s, 8VP101 and 9VP19, the viability of lung cancer cells is inhibited in a concentration dependent manner with IC_50_ values ranging from 6-10 µM (Supp. Fig. 9C). In summary, non-covalent pan-TXNRD inhibitors that bind to the doorstop pocket of these enzymes exhibit significant and broad anti-cancer activity, driven by RNR dysfunction rather than the pro-oxidant mechanism typically associated with traditional inhibitors

## Discussion

Targeting TXNRD1, and to a lesser extent TXNRD2, for cancer therapy is well supported by multiple preclinical studies [6]. The highly reactive selenocysteine residue in the active site of TXNRD1 and TXNRD2 renders these enzymes susceptible to electrophilic attack from compounds such as AF and TRi-1. Covalent modification of TXNRD1 by these agents has been shown to convert TXNRD1 into a selenium compromised thioredoxin reductase-derived apoptotic proteins (secTRAPs) by conferring a gain-of-function NADPH oxidase pro-oxidant activity that promotes cell death [23]. Notably, this pro-oxidant cytotoxicity is often more pronounced with pharmacological inhibition than with siRNA-mediated knockdown of TXNRD1 [24, 25]. Unlike traditional covalent TXNRD inhibitors, our first-in-class non-covalent TXNRD(i)s target a distinct allosteric site known as the *doorstop pocket* [11, 12]. Although treatment with TXNRD(i)s increases intracellular ROS levels, our findings indicate that ROS are not the primary mediators of cytotoxicity in TNBC cells, as antioxidant fails to reverse the observed cell death. In contrast, AF-induced cell death in TNBC is ROS dependent [26]. Similar ROS dependent mechanisms of AF have been reported in other cancers including high grade serious ovarian cancer [27], gastric cancer [28], mesothelioma [29], lung cancer [30, 31], acute lymphoblastic leukemia [32], and B-cell lymphomas [33]. However, it is important to note that the mode of AF-induced cell death varies significantly across cancer types and is influenced by factors such as drug concentration, cancer cell type, and genetic background[34].

Treatment of TNBC cells with TXNRD(i)s activated programs involving several key transcription factors, including NFκB, p21, and p53. This is consistent with the established role of the Trx system in modulating transcription factor activity, as Trx influences the DNA-binding capacity and regulatory functions of NFκB, p21, and p53[6]. Our data further suggest that TXNRD(i)s induce endoplasmic reticulum (ER) stress and activate the UPR, specifically through the PERK– ATF4–CHOP axis. This aligns with previous findings that cytosolic TXNRD1 is required for reducing non-native disulfide bonds in proteins entering the ER [35]. Under prolonged stress conditions, activation of this pathway promotes apoptotic signaling and upregulation of autophagy-related genes [36]. The induction of autophagy likely serves as a compensatory mechanism to degrade misfolded proteins and alleviate proteotoxic stress, which may explain the enrichment of catabolic processes identified by Metascape analysis. However, the precise contribution of UPR activation to TXNRD(i)-mediated cytotoxicity remains unclear and requires further investigation. These findings also suggest that, in addition to apoptosis, autophagy may represent an alternative mechanism of cell death induced by these inhibitors. Notably, AF has been shown to induce paraptosis in breast cancer cells through dual inhibition of TXNRD1 and the proteasome [34]. Paraptosis is a caspase-independent form of programmed cell death characterized by cytoplasmic vacuolation and swelling of the ER and mitochondria. It is typically triggered by sustained ER stress, particularly when the accumulation of misfolded proteins overwhelms the ER’s folding capacity [48]. Given that ER stress emerged as a dominant signature in our transcriptomic analysis, it will be important to determine whether TXNRD(i)s also elicit paraptosis.

Contrary to the anticipated pro-oxidant mechanism involving ROS-mediated cytotoxicity, the anticancer activity of these TXNRD(i)s appears to be primarily driven by disruption of RNR function, as evidenced by a pronounced replication arrest phenotype. We found that exogenous supplementation with dNTPs, the direct substrates of RNR, fully rescued the cell cycle arrest and partially mitigated cell death. The induction of cell death following TXNRD(i) treatment is likely mediated, at least in part, by DNA damage, as indicated by the accumulation of the DNA damage marker γH2AX, which was partially reduced by dNTP supplementation. These findings suggest that impaired RNR activity leads to replication stress and genotoxicity, ultimately activating apoptotic cell death pathways. Supporting the role of RNR dysfunction, we observed additive effects when TXNRD(i)s were combined with standard-of-care DNA-damaging agents used in TNBC treatment. This suggests that co-treatment with TXNRD(i)s may enhance therapeutic efficacy and potentially allow for reduced chemotherapy dosing, thereby improving patient outcomes and quality of life. Notably, TXNRD(i)s synergized with PARP inhibitors, possibly by exacerbating DNA repair deficiencies. These findings further support the rationale for a combination strategy that concurrently targets both DNA damage and repair pathways in TNBC.

RNR requires dithiol electron donors to maintain its catalytic activity. In *E. coli*. RNR both Trx and glutaredoxin (Grx) serve this function [8]. Grx is a small redox enzyme that facilitates thiol-disulfide exchange reactions in the presence of GSH, GSR and NADPH [37]. In human lung cancer cells, high doses of AF induced oxidation of RRM1 leading to dNTP depletion and impaired replication fork elongation [38]. This phenotype was not rescued by co-treatment with the NAC, but was significantly mitigated by supplementation with exogenous nucleotides. Similarly, genetic depletion of TXNRD1 results in RNR oxidation and impaired function in lung cancer cells [38]. In mouse models, conditional deletion of TXNRD1 in T cells leads to defective RNR catalysis and impaired T cell expansion [39]. These findings highlight the critical role of the Trx–TXNRD axis in maintaining mammalian RNR function. Notably, in lung cancers, frequent deletion of the chromosome 8p12 locus, which contains the GSR gene [40], suggests that loss of the GSH–Grx axis may force these cancers to rely more heavily on the Trx–TXNRD pathway. Our data now extend these observations to TNBC, providing the first evidence that the Trx–TXNRD axis is similarly essential for sustaining RNR activity and cellular proliferation in TNBC cells. The mechanistic basis for this reliance in TNBC warrants further investigation. One possibility is that TXNRD(i)s leads to direct oxidation of RRM1, analogous to what has been observed in lung cancer models treated with AF or depletion of TXNRD1 [37]. Alternatively, transcriptional regulation of RNR subunits may play a role. Transcriptomic profiling of TNBC cells treated with TXNRD(i)s revealed significant downregulation of multiple E2F family transcription factors. Given that the promoters of both RRM1 and RRM2 harbor E2F-binding sites [46], suppression of E2F activity may contribute to reduced RNR gene expression. Further studies are needed to elucidate how these transcriptional and post-translational mechanisms converge to regulate RNR function downstream of the Trx–TXNRD axis in TNBC.

Although TXNRD(i)s exhibit broad activity against TNBC, whether specific TNBC subtypes display heightened sensitivity remains to be determined. Transcriptomic analysis revealed modulation of androgen receptor (AR) signaling, suggesting that the luminal androgen receptor (LAR) subtype may derive added benefit from TXNRD(i) treatment. The LAR subtype is characterized by elevated AR expression and has been associated with a poor pathological complete response compared to other TNBC subtypes in patients receiving neoadjuvant chemotherapy [47]. Whether TXNRD(i)s exert enhanced anticancer effects in LAR TNBC warrants further investigation. Given the essential role of RNR in DNA synthesis and repair, it has long been considered a compelling target for cancer therapy [41]. Current clinically approved RNR inhibitors include nucleoside analogs such as gemcitabine and clofarabine, which target the RRM1 subunit, and the radical scavenger hydroxyurea, which inhibits RRM2. However, these agents are limited by significant drawbacks: nucleoside analogs often exhibit dose-limiting toxicity and a narrow therapeutic window, while radical scavengers lack both target specificity and sustained efficacy [41]. Our novel TXNRD(i)s may provide an alternative strategy to disrupt RNR activity, particularly in tumors such as TNBC that exhibit elevated RNR subunit expression and depend on the Trx–TXNRD system.

TXNRD1 is commonly presumed to be the primary therapeutic target in cancer, including TNBC. However, our data suggest that TXNRD2 may be equally relevant, as supported by both genetic and pharmacological approaches. We found that TXNRD(i)s effectively inhibit both cytosolic (TXNRD1) and mitochondrial (TXNRD2) pools of TXNRD, consistent with the prediction that the allosteric doorstop pocket is also present in TXNRD2. Notably, siRNA-mediated knockdown of TXNRD2 induced a robust G1 cell cycle arrest, closely resembling the phenotype observed following TXNRD(i) treatment. This raises the intriguing possibility that TXNRD2 plays a direct role in regulating cell cycle progression. Further studies are warranted to elucidate the precise function of TXNRD2 and to evaluate its potential as a therapeutic target in TNBC, as it may represent an underappreciated vulnerability. In addition to their anticancer activity, our TXNRD(i)s serve as valuable pharmacological tools for advancing our understanding of TXNRD biology. Both TXNRD1 and TXNRD2 are embryonically essential, and shRNA-based depletion of TXNRDs in cancer cells often triggers compensatory upregulation of alternative antioxidant pathways [42]. Our approach offers a unique pharmacological opportunity to dissect TXNRD function and uncover new tumor biology related to TXNRD inhibition in TNBC.

In conclusion, this study demonstrates that our novel, first-in-class non-covalent TXNRD(i)s exhibit potent anticancer activity in TNBC cells through mechanisms distinct from those of traditional TXNRD inhibitors. Our TXNRD(i)s do not rely on pro-oxidant or ROS-dependent mechanisms to exert their effects. Instead, our data support a model in which their primary mechanism of action involves disruption of RNR function, leading to reduced proliferation and induction of DNA damage–mediated cell death. This mechanistic insight has important implications: (i) it can inform the design of next-generation inhibitors with improved anticancer efficacy and safety profiles, and (ii) it offers the potential to develop therapeutic biomarkers for identifying TNBC patients most likely to benefit from TXNRD(i) treatment.

## Figure Legends

**Supp Fig 1.**
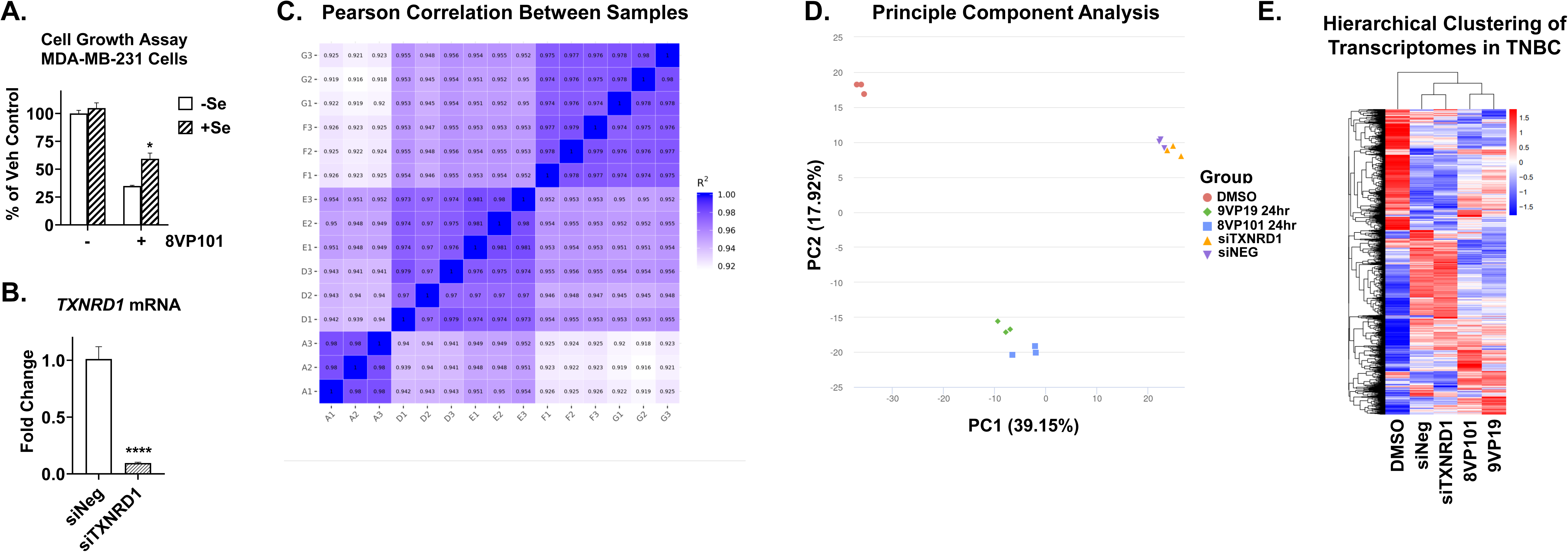
Selenium effect and biological replicates of RNAseq samples. A. MDA-MB-231 cells were pre-treated for 2 hours with 100µM sodium selenite then treated with 2µM TXNRD(i) or vehicle (n = 3). After 72 hours, viability measured was measured with crystal violet. B. Gene expression measured by RT-qPCR in MDA-MB-231 cells transfected with siNEG or siTXNRD1, 10nM each, for 48 hours. Relative gene expression is calculated using the ΔΔCt method normalized to β-actin. C. Pearson correlation analysis of gene expression levels (FPMK) across biological replicates: vehicle (A1-3), 8VP101 (D1-3), 9VP19 (E1-3), siNEG (F1-3), siTXNRD1 (G1-3). D. Principal component analysis of gene expression values (FPKM) across all samples. E. Heatmap showing hierarchical clustering of sample groups (n = 3) based on log₂(FPKM + 1) expression values. Red indicates high expression; blue indicates low expression. Data is presented as the mean +/-sem. Statistical significance was determined by unpaired t-test. *p≤0.05; ****p≤0.0001.

**Supp Fig 2.**
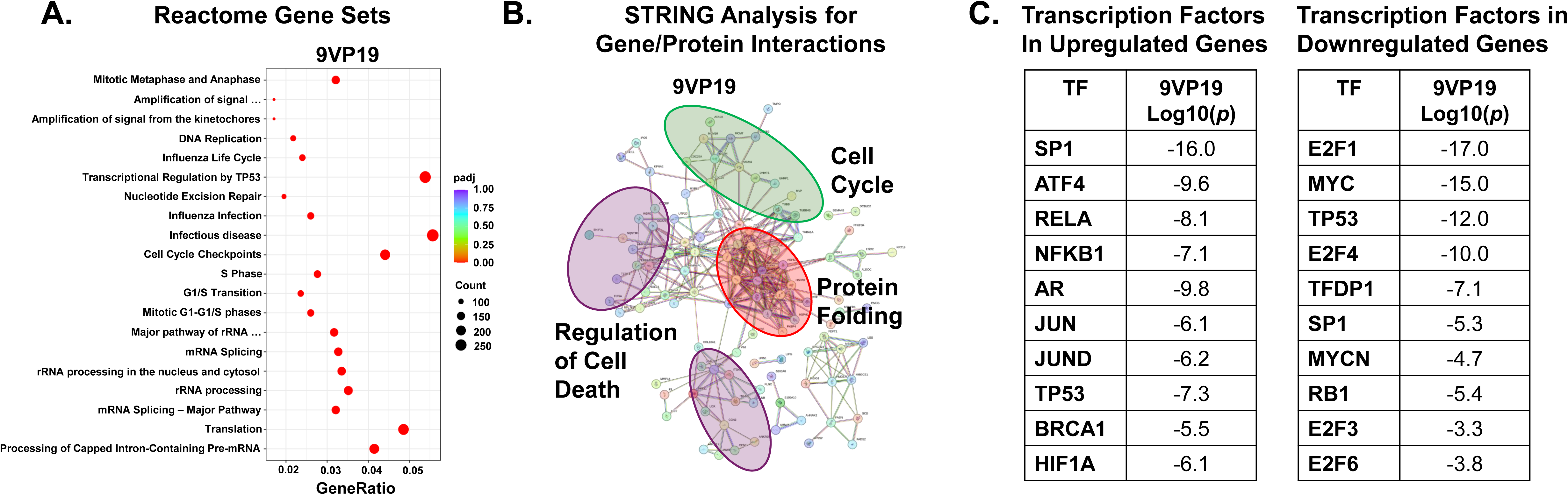
Transcriptomic and bioinformatic analysis induced by 9VP19. A. Reactome enrichment analysis scatter plot. Gene ratio indicates the ratio of the number of differentially expressed genes within the reactome ID (identified by the pathway description) to the number of differentially expressed genes within the reactome database. Count indicates the number of genes within the indicated reactome ID. B. String analysis of the top 150 genes regulated by 9VP19 treatment relative to vehicle. 9VP19: Regulation of Cell Death FDR: 1.37e-05, Protein Folding FDR: 3.39e-07, Cell Cycle FDR: 0.0276. C. Metascape analysis identifying transcription factors regulating the top 2,500 upregulated genes (left) and top 2,500 downregulated genes (right), ranked by –log₁₀(p-value). (–) indicates not detected.

**Supp Fig 3.**
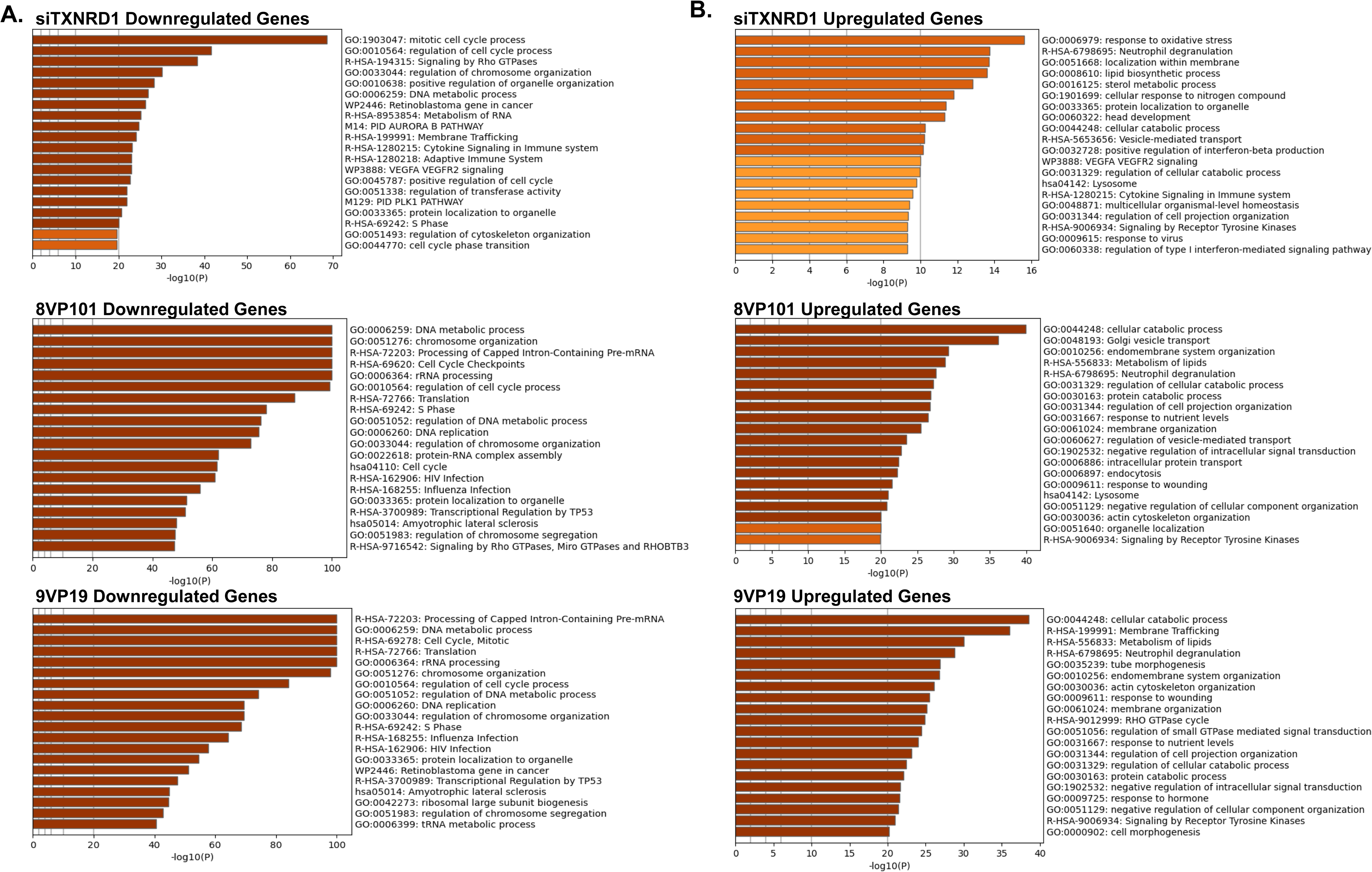
Top pathways identified from Metascape enrichment analysis. The top 2,500 significantly differentially expressed genes (determined by adjusted p < 0.05) were analyzed using Metascape to identify enriched pathways. (A) Downregulated genes and (B) upregulated genes were assessed separately to identify pathways regulated by TXNRD inhibitors (8VP101 and 9VP19) and siTXNRD1. 8VP101 and 9VP19 samples were compared to vehicle-treated controls, while siTXNRD1-transfected samples were compared to siNEG-transfected controls. A. Top significantly downregulated pathways in siTXNRD1 (top), 8VP101 (middle), and 9VP19 (bottom). B. Top significantly upregulated pathways in siTXNRD1 (top), 8VP101 (middle), and 9VP19 (bottom). Pathways are shown ranked according to -log10(p-value).

**Supp Fig 4.**
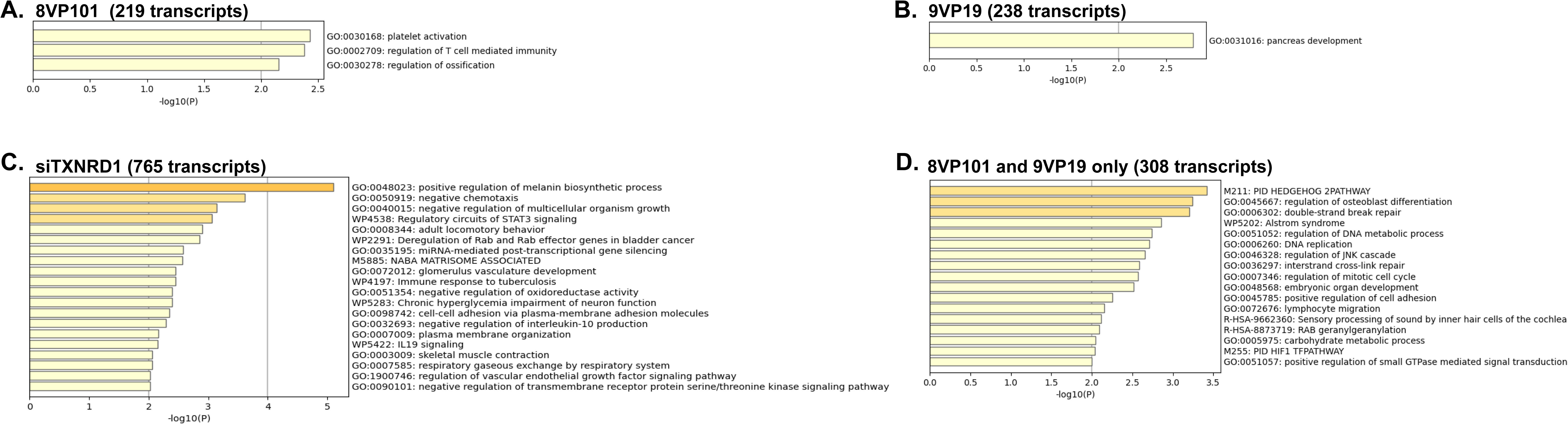
Top pathways from Metascape enrichment analysis of unique non-overlapping genes. Transcripts uniquely regulated by each treatment condition—8VP101, 9VP19, both 8VP101 and 9VP19, and siTXNRD1—were subjected to Metascape enrichment analysis. (A) transcripts (219) uniquely regulated by 8VP101. (B) transcripts (238) uniquely regulated by 9VP19. (C) transcripts (308) uniquely regulated by both 8VP101 and 9VP19. (D) transcripts (765) uniquely regulated by siTXNRD1. Pathways are shown ranked according to -log10(p-value).

**Supp Fig 5.**
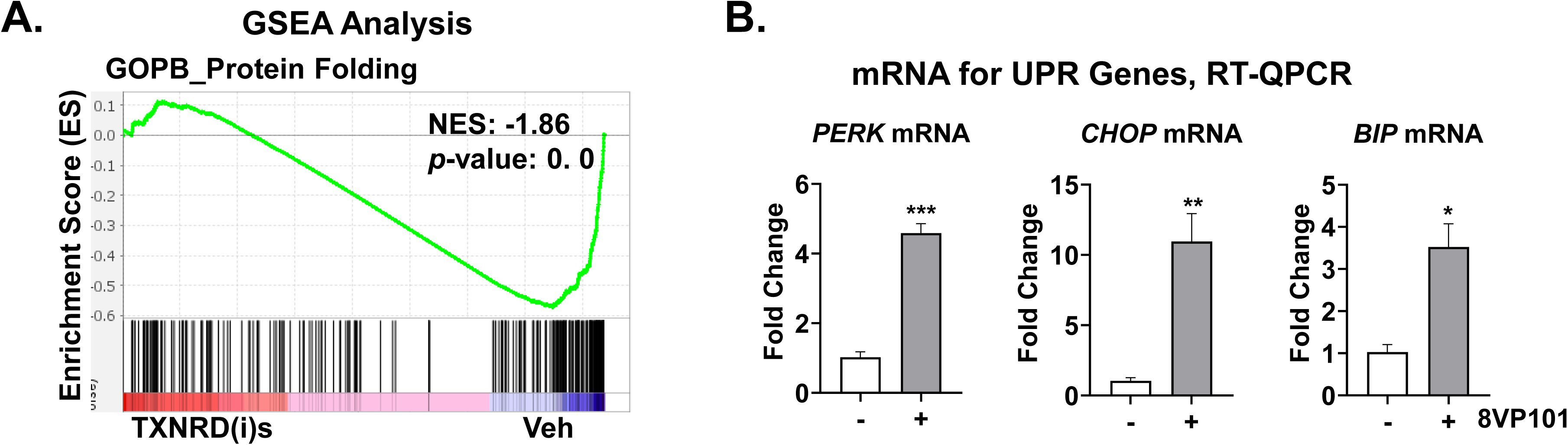
Protein folding signature and UPR genes induced by TXNRD(i)s. A. Gene set enrichment analysis of *Protein Folding* gene set by Gene Ontology Biological Processes. NES and nominal p-value were determined by the GSEA software. B. Gene expression measured by RT-qPCR in MDA-MB-231 cells treated with 10μM 8VP101 (n=3) for 24 hours. Relative gene expression is calculated using the ΔΔCt method normalized to β-actin. Data is presented as the mean +/-sem. P-values were calculated by unpaired t-test. *p≤0.05; **p≤0.005; ***p≤0.001.

**Supp Fig 6.**
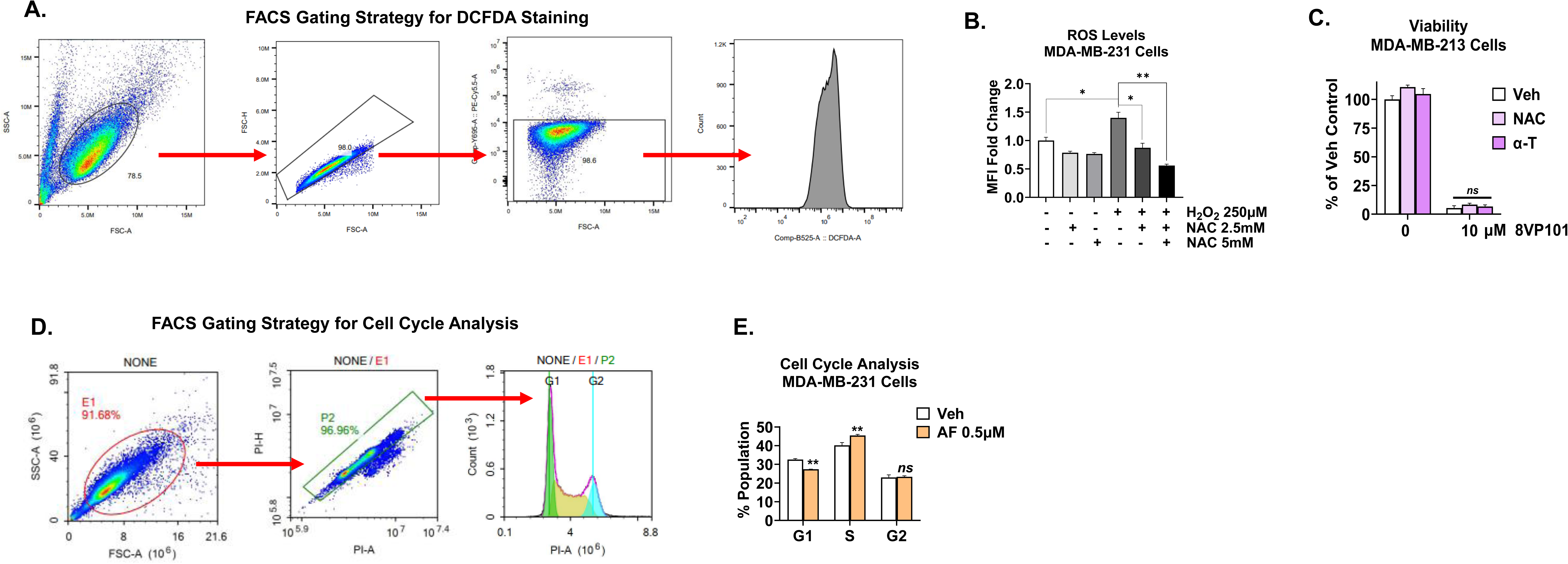
FACS gating strategies, impact of antioxidants and cell cycle analysis induced by AF. A. Representative gating strategy for ROS detection using H_2_DCFDA by flow cytometry. B. ROS quantification using H_2_DCFDA by flow cytometry of MDA-MB-231 cells. Cells were treated with 2.5mM or 5mM N-Acetyl cysteine (NAC) and 250µM H_2_O_2_ for 24 hours, harvested and incubated with 1µM H_2_DCFDA for 15 minutes. Statistical significance was determined by unpaired t-test. C. Cell Viability determined by crystal violet. MDA-MB-231 cells were treated with 10µM 8VP101 and 5mM NAC or 10µM ⍺-tocopherol (⍺-T) for 24 hours. Cells were fixed and stained with crystal violet. Statistical significance was determined by 1-way ANOVA followed by Tukey’s test. D. Representative gating strategy for cell cycle analysis using PI by flow cytometry. E. Cell cycle analysis of MDA-MB-231 cells treated with 0.5μM Auranofin (AF) for 24 hours. Cells were fixed and stained with PI before analysis by flow cytometry. Statistical significance was determined by unpaired t-test. Data is presented as the mean +/-sem. *p≤0.05; **p≤0.005; ns: not significant

**Supp Fig 7.**
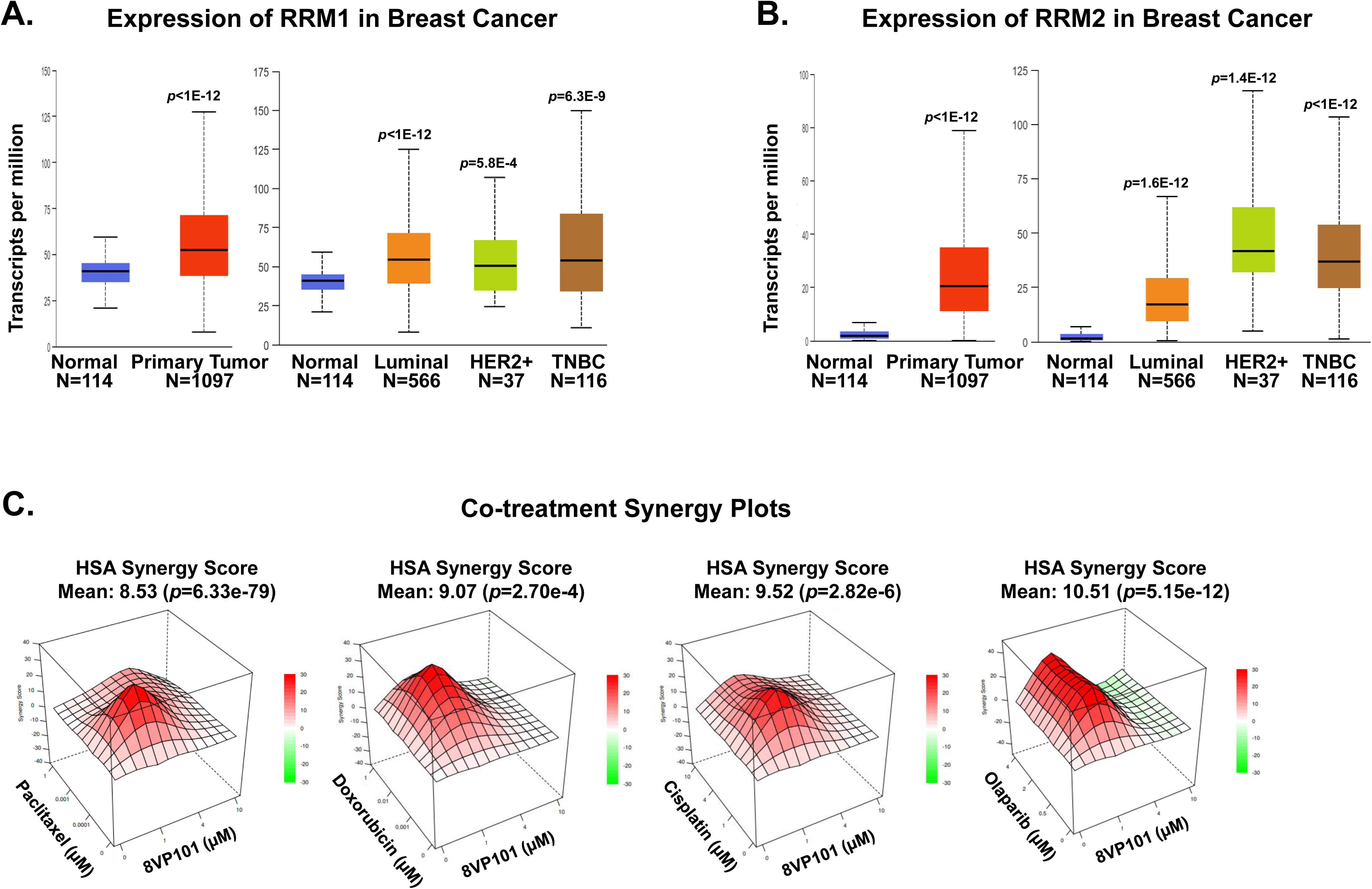
Expression of RNR subunits is elevated in TNBC patient tumors and synergy plots for combination therapies of TXNRD(i) with paclitaxel, doxorubicin, cisplatin, and Olaparib. (A-B) mRNA expression of RRM1 (A) and RRM2 (B) stratified according to breast cancer subtype from TCGA breast cancer database are consistently high TNBC. P-values show significance compared to adjacent normal tissue. C. MDA-MB-231 cells were co-treated with 8VP101 (1µM, 4µM, or 10µM) and a chemotherapeutic, paclitaxel (far left; 0.1nM, 1nM, or 1µM), doxorubicin (left middle; 1M, 10nM, or 1µM), cisplatin (right middle; 1µM, 4µM, or 10µM), or olaparib (far right; 0.5µM, 2µM, or 4µM). Cell viability was quantified after 72 hours. Synergy scores and p-values for percent viability were calculated using SynergyFinder.

**Supp Fig 8.**
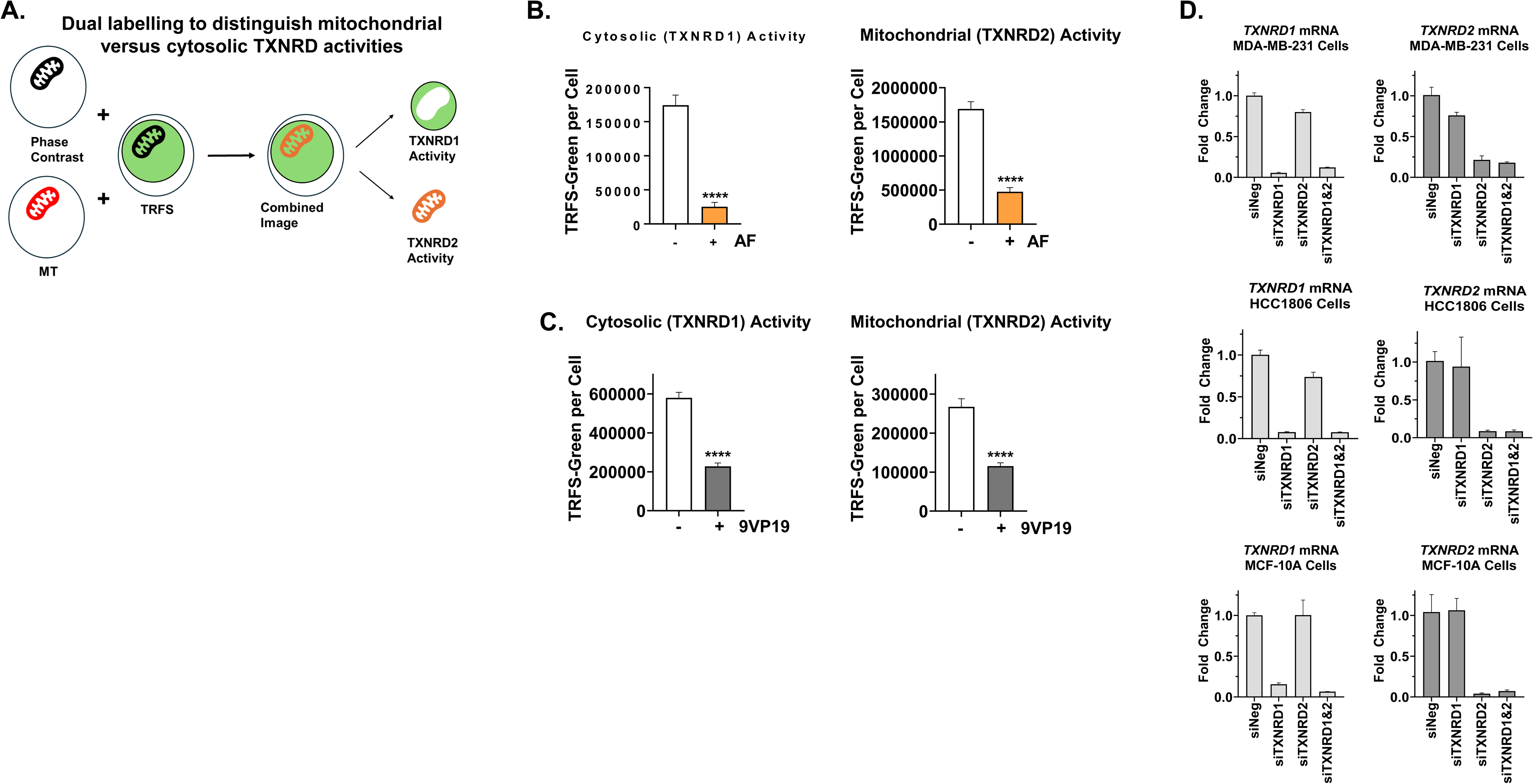
Microscopy-based methodology for determining TXNRD1 versus TXNRD2 activity in live TNBC cells. A. Dual staining methodology scheme. Cells are stained with both TRFS-green for 4 hours and MitoTracker-red for 30 mins and imaged on the Nikon Ti2E inverted microscope at 20x. Colocalization of TRFS-green and MitoTracker-red signal appearing as orange is indictive of mitochondrial or TXNRD2 activity, whereas TRFS-green only is indictive of cytosolic or TXNRD1 activity. B-C. HCC1806 cells are treated with either vehicle control or 0.1µM AF treatment (B) or 4µM 9VP101 (C), then stained and imaged as in (A). Average TRFS-green or MitoTracker-red intensity per cell per field basis was quantified from n=3 biological replicates and n=9 technical replicates. **** p<0.0001. (D) MDA-MB-231, HCC1806 and MCF-10A cells were transfected with siNEG, siTXNRD1, siTXNRD2, or both siTXNRD1 and siTXNRD2, 10nM each, and harvested after 48 hours. Gene expression measured by RT-qPCR.

**Supp Fig 9.**
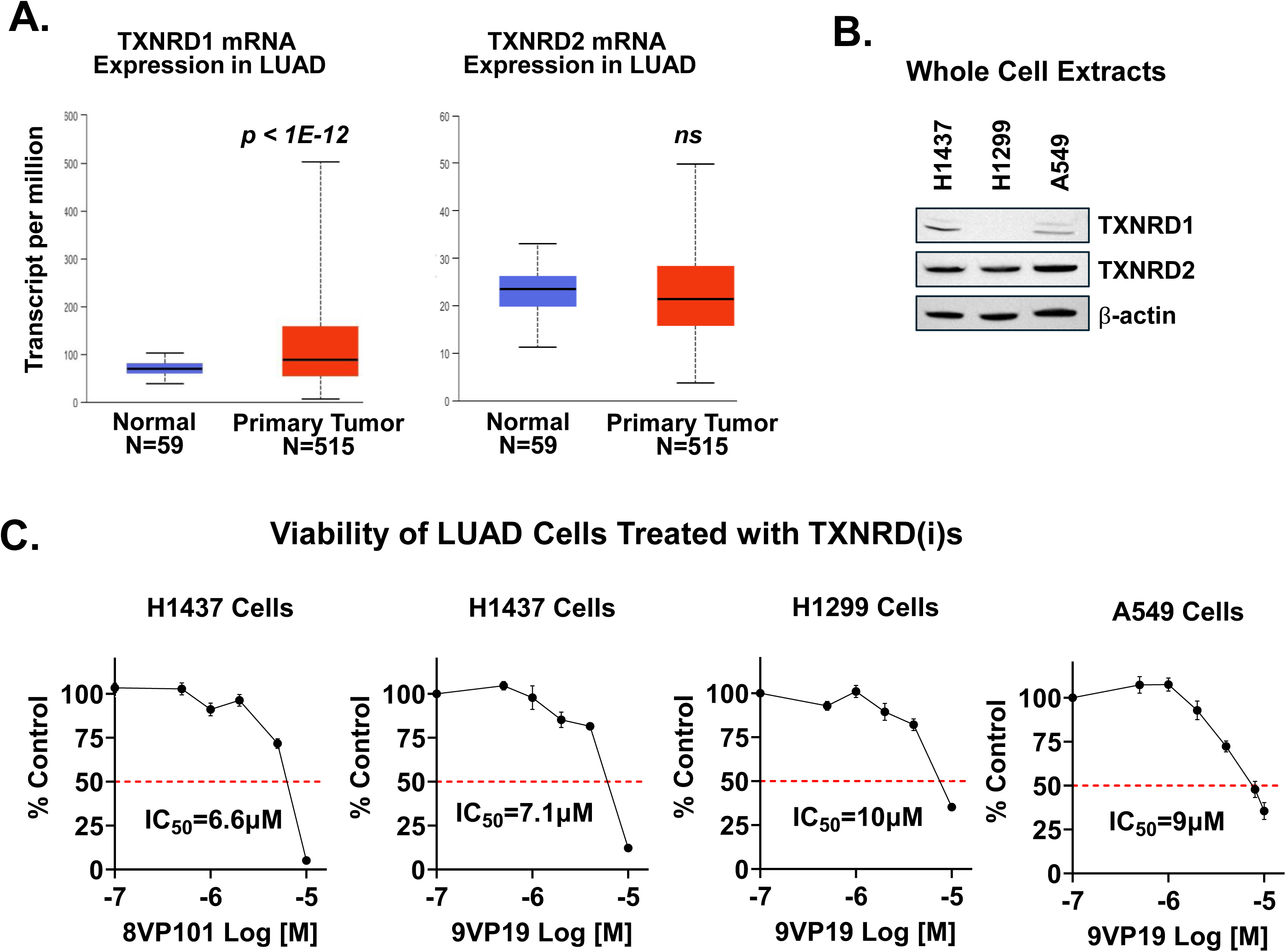
TXNRD(i)s inhibit lung cancer cells. A. RNA expression of TXNRD1 and TXNRD2 from TCGA in lung adenocarcinoma LUAD. B. Protein expression of TXNRD1 and TXNRD2 by western blot analysis in lung cancer cell lines H1437, H1299, and A549. β-actin was used as the loading control. C. Cell viability determined by crystal violet. Lung cancer cells H1437, H1299, and A549 were treated with various concentrations of 8VP101 or 9VP19 for 72 hours. Cell viability was quantified and normalized to vehicle control set to 100 %. The inhibitory concentrations at 50 % (IC_50_) were determined using GraphPad.

## Materials and Methods

### Reagents

The TXNRD inhibitors, 8VP101 and 9VP19, were synthesized as previously reported [12]. Auranofin (AF, A6733) was purchased from Millipore Sigma. Propidium iodide (PI, P1304MP) and PureLinkTM RNase A (12091-021) were purchased from Invitrogen. 2,7-dichloroluorescein diacetate (H2DCFDA, D6883) and N-acetyl cysteine (NAC, A7250) were purchased from Sigma Aldrich. 7-Aminoactinomycin D (7-AAD, A1310) and dNTPs (10297018) were purchased from ThermoFisher Scientific. α-tocopherol (HY-16686) was purchased from MedChemExpress. Silencer® Select Negative Control #2 siRNA (4390846), TXNRD1 Silencer Select Validated siRNA (4427038) and TXNRD2 Silencer Select Validated siRNA (s20782) were purchased from ThermoFisher Scientific. Lipofectamine RNAiMAX Reagent (13778-075) and Opti-MEM (11058-021) were purchased from Fisher Scientific. TRFS-green (HY-115640) was purchased from MedChemExpress. MitoTracker-Red CMXROS (9082P) was purchased from Cell Signaling.

### Cell Lines, Culture Conditions and Treatments

MDA-MB-231 cells were obtained from Dr. Clodia Osipo (Loyola University Chicago) and were maintained in RPMI 1640 medium (Gibco, 11875-093) supplemented with 5% fetal bovine serum (FBS), 2mmol/L L-glutamine, 1% antibiotics penicillin-streptomycin, and 1% non-essential amino acids. HCC1806 cells were obtained from ATCC and maintained in RPMI 1640 medium with HEPES and sodium pyruvate (ATCC, 30-2001) supplemented with 10% FBS, 2mmol/L L-glutamine, 1% antibiotics penicillin-streptomycin, and 1% non-essential amino acids. MCF-10A cells were obtained from ATCC and maintained in DMEM-F12 medium (Gibco, 11330032) supplemented with 10% fetal bovine serum, 20ng/mL epithelial growth factor, 0.5mg/mL hydrocortisone, 100ng/mL cholera toxin, and 10µg/mL insulin. All cells were maintained at 37°C in a humidified atmosphere with 5% CO_2_. H1437, H1299, and A549 cells were obtained from Dr. Maurizio Bocchetta (Loyola University Chicago) and were maintained in RPMI 1640 medium supplemented with 10% FBS. All cell lines were tested for mycoplasma using the LookOut PCR detection kit (Sigma) and validated by STR through ATCC. Inhibitors were dissolved in dimethyl sulfoxide (DMSO) and the final culture volume of inhibitors or vehicle was less than or equal to 0.1%. siRNA transfections were performed according to the manufacturer’s instructions.

### RNASequencing and Bioinformatic Analysis

Total RNA was extracted using TRIzol according to the manufacturer’s instructions. RNA quantity and quality were assessed using a NanoDrop spectrophotometer. RNA-seq library preparation, sequencing, and initial quality control were performed by Novogene. Libraries were sequenced using the Illumina NovaSeq platform employing a paired-end 150bp sequencing strategy. Downstream analysis was conducted using the DESeq2 R package (1.20.0) for differential expression analysis. Gene set enrichment analysis (GSEA) was conducted using the GSEA desktop application (version 4.3.3, Broad Institute). The full gene expression dataset, quantified as FPKM (Fragments Per Kilobase of transcript per Million mapped reads), was used as input. Analysis was performed using the standard (non-pre-ranked) mode with 1,000 phenotype permutations to assess statistical significance. Functional enrichment analysis was performed using Metascape (https://www.metascape.org/) to identify biological pathways and regulatory networks associated with transcriptional changes following TXNRD inhibition. Genes submitted for analysis is specified in figure legend. Default parameters were used: a minimum gene overlap of 3, a p-value cutoff of 0.01, and an enrichment factor threshold of 1.5. Results were visualized using Metascape’s clustering and pathway annotation tools.

### Cell Viability

Cells were seeded in 24-well plates with fresh media. After overnight attachment, cells were treated with the indicated TXNRD1 inhibitors for 72 hours. Cells were stained with 1% crystal violet in methanol and water (1:4), solubilized in 1% sodium dodecyl sulfate (SDS) and the absorbance was measured at 570nm using the BioTek Synergy H1 Plate Reader.

### Flow Cytometry

Detection of ROS was measured using H_2_DCFDA. H_2_DCFDA was used as previously describe in the literature [43]. Cells were harvested by trypsinization and loaded with 1μM H_2_DCFDA in serum free media for 15 minutes at 37°C. Cells were then washed twice with ice-cold PBS, resuspended in ice-cold PBS with 3µg/mL of 7-AAD and analyzed immediately using NovoCyte Quanteon Flow Cytometer by Agilent Technologies. Mean fluorescence intensity (MFI) was used as a measure of ROS and data were analyzed with Flowjo v11. For 5-ethynyl-2′-deoxyuridine (EdU) incorporation, cells were stained according to the manufacturer’s instructions using an EdU Staining Proliferation Kit iFluor 488 (ab219801) from Abcam. Proliferation was immediately analyzed using NovoCyte Quanteon Flow Cytometer by Agilent Technologies. For cell cycle analysis, cells were collected using trypsin and washed twice with ice-cold PBS. Cells were resuspended in 0.5mL of ice-cold PBS and 1mL of 100% Ethanol was added in dropwise while vortexing, followed by incubation for 1 hour at 4°C. Cells were then resuspended in RNase A (amount) and PI (amount) for 1 hour at room temperature, then immediately analyzed using NovoCyte Quanteon Flow Cytometer by Agilent Technologies. Gating strategy outlined in appendix A (Figure 27C). Cell death was measured using AlexaFluor^TM^ 488 Annexin V/Dead Cell Apoptosis Kit (V13241) from ThermoFisher Scienticifc according to the manufacturer’s instructions. Apoptosis was immediately analyzed using NovoCyte Quanteon Flow Cytometer by Agilent Technologies.

### Microscopy in Live Cells

Cells were seeded at high confluency in a 24-well plate. The next day, cells were pretreated with the inhibitors for 2 hours. Cells were then incubated with 10µM TRFS-green at 37°C for 4 hours followed by 100nM MitoTracker-red CMXRos (CellSignaling Technology, cat# 9082) for 30 mins and imaged on the Nikon Ti2E inverted microscope at 20x using the GFP/FITC filter cube with excitation of 480nm for TRFS-green and TRITC/CY3 filter cube with excitation of 540nm for MitoTracker-red. Utilizing the NIS-Element application, regions of interest (ROIs) were drawn around the perimeter of each cell in each combined image’s field. TRFS-green and MitoTracker-red intensity were threshold and used as reference binaries for intersection (TXNRD2 activity pool) and subtraction (TXNRD1 activity pool) binaries. Average intensity per cell per field was calculated from the total SumGreen intensity divided by total cell per field for TRFS-green, intersection and subtraction binaries.

### Western Blot

Whole cell extracts were prepared using the M-PER reagent (ThermoFisher Scientific). Proteins were separated by SDS-PAGE (Invitrogen), transferred to nitrocellulose membranes using an iBlot 2 instrument (Invitrogen), blocked for 1 hour in TBS/T buffer containing 5 % non-fat dry milk and incubated with the indicated primary antibody (diluted in 2.5 % milk in TBS/T) overnight. The next day, membranes were incubated with secondary antibody for 1 hour and the signal visualized on the iBright CL1000 Imaging System (Invitrogen) using the Pierce Supersignal West Pico chemiluminescent substrate (ThermoFisher Scientific). The antibody for β-actin (A5441) was purchased from Sigma Aldrich. The antibody for γ-H2AX (2577S) was purchased from Cell Signaling. The antibody for p21 (10355-1-AP) was purchased from ProteinTech. The antibodies for TXNRD1 (15140S) and TXNRD2 (12029S) were purchased from Cell Signaling.

### RT-Quantitative PCR (QPCR)

Total RNA was isolated using TRIzol according to manufacturer’s instructions. RNA (0.5µg) was reverse transcribed in a total volume of 10μL using 200U of M-MLV reverse transcriptase, 100ng random hexamer, 0.5mM deoxy-NTP and 10mM DTT. The resulting cDNA was mixed with SYBR Green Master mix, forward and reverse primers and the amplifications were performed using a QuantStudio3 instrument (ThermoFisher Scientific) according to manufacturer’s instructions. Fold change was calculated using the ΔΔCt method with β-actin serving as the internal control. All QPCR primers used were validated and are available upon request.

### In vivo Studies

Mouse experiments were performed at the Loyola University Chicago animal facility and conducted in accordance with institutional procedures and guidelines after prior approval from the Institutional Animal Care and Use Committee (IACUC). Female athymic nude mice (nu/nu), aged 5-weeks-old, were purchased from Envigo and allowed to acclimate for one week. One or two million MDA-MB-231 or HCC1806 cells, respectively in 100µL PBS were bilaterally injected orthotopically into the thoracic mammary glands. Tumor formation was monitored by palpation and once tumors were detected (∼50mm^3^ in size), mice were randomized into either vehicle control (10% DMSO, 10% tween-80, and 80% PBS for 100µL volume) or treatment groups of 8VP101, 50mg/kg. Mice received daily treatments via intraperitoneal (IP) injection. Tumor sizes were measured daily with an electronic caliper. The tumor volume was calculated as length/2 × width^2^ × π. Tumor were excised and processed for gene expression.

### Statistical Analysis

Experiments were done in biological triplicate unless otherwise stated. Data are presented as mean ± sem from at least three independent determinations. Statistical analysis consisted of 1-way ANOVA followed by Tukey posttest, unpaired t-test, and Kruskal Wallis test followed by Dunns posttest, as appropriate.

## Acknowledgment

The study was funded in part by a pilot grant to I.K. from the LUC Cardinal Bernardin Cancer Center, the Institute for Translational Medicine pilot award to I.K., and P.A.P., the US NIH/NIAID R01AI177493 to P.A.P., D.L.W. and F.A., and by the UIC Proof-of-Concept study to P.A.P., the University of Illinois Collaborative Engagement in Novel Therapeutic Research and Enterprise: UICentre, We acknowledge OpenEye/Cadence for providing P.A.P. laboratory with an academic license for the software used in these studies.

## References

[1] D. Hanahan, R.A. Weinberg, Hallmarks of cancer: the next generation, Cell 144(5) (2011) 646–74.

[2] F. Xing, Q. Hu, Y. Qin, J. Xu, B. Zhang, X. Yu, W. Wang, The Relationship of Redox With Hallmarks of Cancer: The Importance of Homeostasis and Context, Front Oncol 12 (2022) 862743.

[3] I.S. Harris, A.E. Treloar, S. Inoue, M. Sasaki, C. Gorrini, K.C. Lee, K.Y. Yung, D. Brenner, C.B. Knobbe-Thomsen, M.A. Cox, A. Elia, T. Berger, D.W. Cescon, A. Adeoye, A. Brustle, S.D. Molyneux, J.M. Mason, W.Y. Li, K. Yamamoto, A. Wakeham, H.K. Berman, R. Khokha, S.J. Done, T.J. Kavanagh, C.W. Lam, T.W. Mak, Glutathione and thioredoxin antioxidant pathways synergize to drive cancer initiation and progression, Cancer Cell 27(2) (2015) 211–22.

[4] E.S. Arner, A. Holmgren, The thioredoxin system in cancer, Semin Cancer Biol 16(6) (2006) 420–6.

[5] E.S. Arner, Focus on mammalian thioredoxin reductases--important selenoproteins with versatile functions, Biochim Biophys Acta 1790(6) (2009) 495–526.

[6] R. Gencheva, E.S.J. Arner, Thioredoxin Reductase Inhibition for Cancer Therapy, Annu Rev Pharmacol Toxicol 62 (2022) 177–196.

[7] R. Gencheva, Q. Cheng, E.S.J. Arner, Thioredoxin reductase selenoproteins from different organisms as potential drug targets for treatment of human diseases, Free Radic Biol Med 190 (2022) 320–338.

[8] R. Sengupta, A. Holmgren, Thioredoxin and glutaredoxin-mediated redox regulation of ribonucleotide reductase, World J Biol Chem 5(1) (2014) 68–74.

[9] T.C. Laurent, E.C. Moore, P. Reichard, Enzymatic Synthesis of Deoxyribonucleotides. Iv. Isolation and Characterization of Thioredoxin, the Hydrogen Donor from Escherichia Coli B, J Biol Chem 239 (1964) 3436–44.

[10] C.B. Prasad, A. Oo, Y. Liu, Z. Qiu, Y. Zhong, N. Li, D. Singh, X. Xin, Y.J. Cho, Z. Li, X. Zhang, C. Yan, Q. Zheng, Q.E. Wang, D. Guo, B. Kim, J. Zhang, The thioredoxin system determines CHK1 inhibitor sensitivity via redox-mediated regulation of ribonucleotide reductase activity, Nat Commun 15(1) (2024) 4667.

[11] M. Ardini, S.Y. Aboagye, V.Z. Petukhova, I. Kastrati, R. Ippoliti, G.R.J. Thatcher, P.A. Petukhov, D.L. Williams, F. Angelucci, The “Doorstop Pocket” In Thioredoxin Reductases horizontal line An Unexpected Druggable Regulator of the Catalytic Machinery, J Med Chem 67(18) (2024) 15947–15967.

[12] V.Z. Petukhova, S.Y. Aboagye, M. Ardini, R.P. Lullo, F. Fata, M.E. Byrne, F. Gabriele, L.M. Martin, L.N.M. Harding, V. Gone, B. Dangi, D.D. Lantvit, D. Nikolic, R. Ippoliti, G. Effantin, W.L. Ling, J.J. Johnson, G.R.J. Thatcher, F. Angelucci, D.L. Williams, P.A. Petukhov, Non-covalent inhibitors of thioredoxin glutathione reductase with schistosomicidal activity in vivo, Nat Commun 14(1) (2023) 3737.

[13] B. Flowers, A. Rullo, A. Zhang, K. Chang, V.Z. Petukhova, S.Y. Aboagye, F. Angelucci, D.L. Williams, S. Kregel, P.A. Petukhov, I. Kastrati, Pleiotropic anti-cancer activities of novel non-covalent thioredoxin reductase inhibitors against triple negative breast cancer, Free Radic Biol Med 227 (2025) 201–209.

[14] R. Gencheva, L. Coppo, E.S.J. Arner, X. Ren, Selenium supplementation protects cancer cells from the oxidative stress and cytotoxicity induced by the combination of ascorbate and menadione sodium bisulfite, Free Radic Biol Med 233 (2025) 317–329.

[15] B. Kalyanaraman, V. Darley-Usmar, K.J. Davies, P.A. Dennery, H.J. Forman, M.B. Grisham, G.E. Mann, K. Moore, L.J. Roberts, 2nd, H. Ischiropoulos, Measuring reactive oxygen and nitrogen species with fluorescent probes: challenges and limitations, Free Radic Biol Med 52(1) (2012) 1–6.

[16] K.R. Atkuri, J.J. Mantovani, L.A. Herzenberg, L.A. Herzenberg, N-Acetylcysteine--a safe antidote for cysteine/glutathione deficiency, Curr Opin Pharmacol 7(4) (2007) 355–9.

[17] J.M. Tucker, D.M. Townsend, Alpha-tocopherol: roles in prevention and therapy of human disease, Biomed Pharmacother 59(7) (2005) 380–7.

[18] A. Sharma, K. Singh, A. Almasan, Histone H2AX phosphorylation: a marker for DNA damage, Methods Mol Biol 920 (2012) 613–26.

[19] A. Ianevski, A.K. Giri, T. Aittokallio, SynergyFinder 3.0: an interactive analysis and consensus interpretation of multi-drug synergies across multiple samples, Nucleic Acids Res 50(W1) (2022) W739–W743.

[20] L. Zhang, D. Duan, Y. Liu, C. Ge, X. Cui, J. Sun, J. Fang, Highly selective off-on fluorescent probe for imaging thioredoxin reductase in living cells, J Am Chem Soc 136(1) (2014) 226–33.

[21] K. Neikirk, A.G. Marshall, B. Kula, N. Smith, S. LeBlanc, A. Hinton, Jr., MitoTracker: A useful tool in need of better alternatives, Eur J Cell Biol 102(4) (2023) 151371.

[22] X. Zhang, K. Selvaraju, A.A. Saei, P. D’Arcy, R.A. Zubarev, E.S. Arner, S. Linder, Repurposing of auranofin: Thioredoxin reductase remains a primary target of the drug, Biochimie 162 (2019) 46–54.

[23] W.C. Stafford, X. Peng, M.H. Olofsson, X. Zhang, D.K. Luci, L. Lu, Q. Cheng, L. Trésaugues, T.S. Dexheimer, N.P. Coussens, M. Augsten, H.M. Ahlzén, O. Orwar, A. Östman, S. Stone-Elander, D.J. Maloney, A. Jadhav, A. Simeonov, S. Linder, E.S.J. Arnér, Irreversible inhibition of cytosolic thioredoxin reductase 1 as a mechanistic basis for anticancer therapy, Sci Transl Med 10(428) (2018).

[24] M.H. Yoo, X.M. Xu, B.A. Carlson, A.D. Patterson, V.N. Gladyshev, D.L. Hatfield, Targeting thioredoxin reductase 1 reduction in cancer cells inhibits self-sufficient growth and DNA replication, PLoS One 2(10) (2007) e1112.

[25] M.-H. Yoo, X.-M. Xu, B.A. Carlson, V.N. Gladyshev, D.L. Hatfield, Thioredoxin Reductase 1 Deficiency Reverses Tumor Phenotype and Tumorigenicity of Lung Carcinoma Cells *, Journal of Biological Chemistry 281(19) (2006) 13005–13008.

[26] O.E. Chepikova, D. Malin, E. Strekalova, E.V. Lukasheva, A.A. Zamyatnin, Jr., V.L. Cryns, Lysine oxidase exposes a dependency on the thioredoxin antioxidant pathway in triple-negative breast cancer cells, Breast Cancer Res Treat 183(3) (2020) 549–564.

[27] F.H. Abdalbari, E. Martinez-Jaramillo, B.N. Forgie, E. Tran, E. Zorychta, A.A. Goyeneche, S. Sabri, C.M. Telleria, Auranofin Induces Lethality Driven by Reactive Oxygen Species in High-Grade Serous Ovarian Cancer Cells, Cancers (Basel) 15(21) (2023).

[28] P. Zou, M. Chen, J. Ji, W. Chen, X. Chen, S. Ying, J. Zhang, Z. Zhang, Z. Liu, S. Yang, G. Liang, Auranofin induces apoptosis by ROS-mediated ER stress and mitochondrial dysfunction and displayed synergistic lethality with piperlongumine in gastric cancer, Oncotarget 6(34) (2015) 36505–21.

[29] B.R. You, W.H. Park, Auranofin induces mesothelioma cell death through oxidative stress and GSH depletion, Oncol Rep 35(1) (2016) 546–551.

[30] X.Y. Cui, S.H. Park, W.H. Park, Anti-Cancer Effects of Auranofin in Human Lung Cancer Cells by Increasing Intracellular ROS Levels and Depleting GSH Levels, Molecules 27(16) (2022).

[31] V. Ehrenfeld, J.R. Heusel, S. Fulda, S.J.L. van Wijk, ATM inhibition enhances Auranofin-induced oxidative stress and cell death in lung cell lines, PLoS One 15(12) (2020) e0244060.

[32] M. Karsa, A. Kosciolek, A. Bongers, A. Mariana, T. Failes, A.J. Gifford, U.R. Kees, L.C. Cheung, R.S. Kotecha, G.M. Arndt, M. Haber, M.D. Norris, R. Sutton, R.B. Lock, M.J. Henderson, K. Somers, Exploiting the reactive oxygen species imbalance in high-risk paediatric acute lymphoblastic leukaemia through auranofin, Br J Cancer 125(1) (2021) 55–64.

[33] J. Wang, J. Wang, E. Lopez, H. Guo, H. Zhang, Y. Liu, Z. Chen, S. Huang, S. Zhou, A. Leeming, R.J. Zhang, D. Jung, H. Shi, H. Grundman, D. Doakes, K. Cui, C. Jiang, M. Ahmed, K. Nomie, B. Fang, M. Wang, Y. Yao, L. Zhang, Repurposing auranofin to treat TP53-mutated or PTEN-deleted refractory B-cell lymphoma, Blood Cancer J 9(12) (2019) 95.

[34] M.J. Seo, I.Y. Kim, D.M. Lee, Y.J. Park, M.-Y. Cho, H.J. Jin, K.S. Choi, Dual inhibition of thioredoxin reductase and proteasome is required for auranofin-induced paraptosis in breast cancer cells, Cell Death & Disease 14(1) (2023) 42.

[35] G.J. Poet, O.B. Oka, M. van Lith, Z. Cao, P.J. Robinson, M.A. Pringle, E.S. Arnér, N.J. Bulleid, Cytosolic thioredoxin reductase 1 is required for correct disulfide formation in the ER, Embo j 36(5) (2017) 693–702.

[36] C. He, D.J. Klionsky, Regulation mechanisms and signaling pathways of autophagy, Annu Rev Genet 43 (2009) 67–93.

[37] C.H. Lillig, C. Berndt, A. Holmgren, Glutaredoxin systems, Biochim Biophys Acta 1780(11) (2008) 1304–17.

[38] C.B. Prasad, A. Oo, Y. Liu, Z. Qiu, Y. Zhong, N. Li, D. Singh, X. Xin, Y.-J. Cho, Z. Li, X. Zhang, C. Yan, Q. Zheng, Q.-E. Wang, D. Guo, B. Kim, J. Zhang, The thioredoxin system determines CHK1 inhibitor sensitivity via redox-mediated regulation of ribonucleotide reductase activity, Nature Communications 15(1) (2024) 4667.

[39] J. Muri, S. Heer, M. Matsushita, L. Pohlmeier, L. Tortola, T. Fuhrer, M. Conrad, N. Zamboni, J. Kisielow, M. Kopf, The thioredoxin-1 system is essential for fueling DNA synthesis during T-cell metabolic reprogramming and proliferation, Nat Commun 9(1) (2018) 1851.

[40] Wistuba, II, C. Behrens, A.K. Virmani, S. Milchgrub, S. Syed, S. Lam, B. Mackay, J.D. Minna, A.F. Gazdar, Allelic losses at chromosome 8p21-23 are early and frequent events in the pathogenesis of lung cancer, Cancer Res 59(8) (1999) 1973–9.

[41] S.E. Huff, J.M. Winter, C.G. Dealwis, Inhibitors of the Cancer Target Ribonucleotide Reductase, Past and Present, Biomolecules 12(6) (2022).

[42] A.A. Bondareva, M.R. Capecchi, S.V. Iverson, Y. Li, N.I. Lopez, O. Lucas, G.F. Merrill, J.R. Prigge, A.M. Siders, M. Wakamiya, S.L. Wallin, E.E. Schmidt, Effects of thioredoxin reductase-1 deletion on embryogenesis and transcriptome, Free Radic Biol Med 43(6) (2007) 911–23.

[43] O.G. Lyublinskaya, J.S. Ivanova, N.A. Pugovkina, I.V. Kozhukharova, Z.V. Kovaleva, A.N. Shatrova, N.D. Aksenov, V.V. Zenin, Y.A. Kaulin, I.A. Gamaley, N.N. Nikolsky, Redox environment in stem and differentiated cells: A quantitative approach, Redox Biol 12 (2017) 758–769.

